# Tumour-derived extracellular vesicles within the therapy-induced senescent secretome distinctly suppress breast cancer via DKK1-mediated inflammatory response

**DOI:** 10.1101/2024.03.29.586905

**Authors:** Matius Robert, Rekha Jakhar, Bijin Veonice Au, Gracie Wee Ling Eng, Meng Wei, Alvin Kunyao Gao, You Heng Chuah, Karishma Sachaphibulkij, Isabelle Bonne, Kah Jing Lim, Indrajit Sinha, Daniel Boon Leng Teh, Lina Hsiu Kim Lim, Prashant Kumar, Navin Kumar Verma, Newman Siu Kwan Sze, Elaine Hsuen Lim, Derrick Sek Tong Ong, Jit Kong Cheong, Koji Itahana, John Edward Connolly, Karen Carmelina Crasta

## Abstract

Triple-negative breast cancers (TNBC), associated with poor prognosis and high tumour recurrence, are often-treated with taxanes in first-line treatment regimens. However, acquired disease resistance can often set in, hampering clinical efficacy. One avenue that could engender therapy resistance is therapy-induced senescence (TIS), as they represent a population of residual disease and are highly secretory. Although it is known that TIS can contribute to tumour development and therapy resistance via the therapy-induced secretome, the underlying molecular mechanisms are not fully understood. In this study, we sought to dissect the role of the TNBC-derived TIS-associated secretome in chemoresponse. We found that paclitaxel-treated cells induced mitotic slippage and entered senescence as tetraploid cells. The therapy-induced SASP was found to be enriched in soluble cytokines and other pro-tumorigenic factors linked to tumour recurrence and distant metastasis. Interestingly, we found that senescence-associated small extracellular vesicles (sEVs) or exosomes, an underappreciated component of SASP, increased genomic instability, ROS and anti-tumour activity. Exosomal proteomic and transcriptomic profiling further revealed DKK1, a negative regulator of WNT signalling, to be enriched in TIS-sEVs. Further investigation demonstrated DKK1-control of inflammatory cytokines production to confer reduced tumour activity in recipient TNBC cancer cells. Taken together, this study revealed unexpected findings where TIS-sEVs confer opposing tumourigenic outcomes to that elicited by TIS-SASP, indicating that sEVs should be considered as distinct SASP entities.

## INTRODUCTION

Although constituting a minority of breast cancers, triple-negative breast cancers (TNBC), the most aggressive breast cancer subtype with poor prognosis, account for disproportionately high metastases, elevated risk of disease relapse and high mortality rates (Yin *et al*, 2020). Since treatment for TNBC is restricted due to lack of specific targets, cytotoxic therapy such as taxane-based chemotherapy, either alone or in combination, remain as mainstay first-line treatment regimen (neoadjuvant or adjuvant) for patients with diagnosis of early-stage, locally advanced and metastatic TNBC (Mustacchi & De Laurentiis, 2015). While most tumours are initially sensitive to treatment, acquired drug resistance can set in, hampering long-term clinical efficacy. Mounting evidence from animal models indicate that cancer therapies can themselves mediate drug resistance via non-cell-autonomous mechanisms, where the altered tumour secretome of drug-sensitive cancer cells confers physiological changes within the tumour microenvironment and supports growth of drug-resistant clones (Keklikoglou *et al*, 2019; Obenauf *et al*, 2015; Shi *et al*, 2014). **This highlights the importance of understanding mechanisms by which therapy-induced tumour secretomes exert paracrine influence for potential early detection of disease relapse and better design of treatment regimens for enhanced efficacy.**

**An important yet often-overlooked tumour secretome-modulating avenue is therapy-induced senescence (TIS).** Cellular senescence, a common therapeutic outcome besides cell death, manifests as a stable cell cycle arrest, conferring tumour suppression. Senescent cells exert their non-cell-autonomous influence on neighbouring cells and the microenvironment via a bioactive secretome consisting of cytokines, chemokines, growth factors and matrix metalloproteinases, termed senescence-associated secretory phenotype (SASP) (Coppe *et al*, 2010; Coppe *et al*, 2008). Soluble SASP factors (sSASP) have been extensively investigated in recent years. In addition to reinforcing senescence in an autocrine manner (Acosta *et al*, 2013; Acosta *et al*, 2008; Kuilman *et al*, 2008), a key role of the sSASP is its paracrine recruitment of immune cells that eliminate senescent cells (Kale *et al*, 2020; Kang *et al*, 2011; Krizhanovsky *et al*, 2008; Xue *et al*, 2007). Interestingly, paracrine action conferred by sSASP can also be pro-tumourigenic, for example, by conferring invasive and migratory properties in neighbouring cells (Bavik *et al*, 2006; Coppe *et al*., 2010; Faget *et al*, 2019; Jakhar *et al*, 2018; Wong *et al*, 2018). Importantly, SASP-driven immunosuppressive microenvironments have also been shown to engender cancer development and tumour recurrence (Matsuda *et al*, 2023; Ruhland *et al*, 2016).

**Although sSASP has been the pivotal focus of SASP function, other entities such as small extracellular vesicles (sEVs), have emerged as important yet under-appreciated components of the senescent secretome (Fafian-Labora & O’Loghlen, 2020; Jakhar & Crasta, 2019). While it is now established that SASP induction pose prognostic and therapeutic implications in cancer progression and therapy response** [well-reviewed in (Faget *et al*., 2019; Schmitt *et al*, 2022)], the role of senescence-associated EVs in this regard remains largely unknown (Misawa *et al*, 2020; Wallis *et al*, 2020). sEVs, such as exosomes, are lipid bilayer- enclosed vesicles (size 30nm -150nm) released from cells that serve as vehicles of transfer for biological macromolecules such as proteins, lipids and nucleic acids, from parental to recipient cells (Colombo *et al*, 2014). In this way, sEVs mediate intercellular communication and influence physiological and pathological functions in recipient cells. Importantly, sEVs can modulate the tumour microenvironment, thereby influencing tumour progression and drug resistance (Clancy & D’Souza-Schorey, 2023; Mathieu *et al*, 2019). Indeed, chemotherapy-elicited EVs have been shown to confer mammary tumour lung metastasis in mice by promoting inflammatory lung endothelial cell activation (Keklikoglou *et al*., 2019). Similar to sSASP, EVs also possess opposing roles and can elicit tumour suppression. For instance, DNA-containing EVs derived from chemotherapy-treated tumours potentiate anti-tumour immunity via cGAS-STING signalling (Kitai *et al*, 2017). One can extrapolate that the immunomodulatory effect of senescent tumour cell-derived EVs alone, in the absence of classical SASP factors such as sSASP, hold potential to influence tumour development and therapy resistance as well. **However, studies have focused almost exclusively on senescent normal and stromal cells with a paucity of insight into the biological impact on chemoresponse by sEVs derived from tumour cells rendered senescent by chemotherapies.** For instance, sEVs secreted from doxorubicin-induced senescent non-transformed cells were shown to promote cancer cell proliferation *in vitro* via the receptor tyrosine kinase EphA2 (Takasugi *et al*, 2017), while more recently, senescent stromal cells were shown to promote chemoresistance via SIRT1 loss-mediated overproduction of sEVs (Han *et al*, 2020).

In this study, we set out to investigate the role of sEVs derived from paclitaxel-induced senescent TNBCs in tumour development. **Strikingly, comparative assessment between SASP (whole secretome consisting of sSASP and sEVs) and senescent tumour cell-derived sEVs alone shortly after senescence establishment uncovered unexpected tumour phenotypic profiles.**

## RESULTS

### SASP factors from paclitaxel-induced senescent TNBC cells elicit tumourigenic effects

Our earlier work demonstrated that human breast tissues with invasive ductal carcinoma post- paclitaxel (PTX) treatment displayed senescence-associated markers at the invasive front (He *et al*, 2018; Jakhar *et al*., 2018). In order to evaluate tumourigenic impact of the secretome of PTX-induced senescent TNBCs, we first established conditions to render cells senescent. To this end, TP53- deficient MDA-MB-231 and TP53-proficient, chromosomally-stable CAL-51 cells were treated with 150 nM PTX for three days to allow mitotically-arrested cells to undergo mitotic slippage and enter therapy-induced senescence (TIS) **(Supplementary Figure S1A)** (Brito & Rieder, 2006; Cheng & Crasta, 2017). Indeed, a substantial number of MDA-MB-231 cells collected three days post-treatment (D3) were viable (∼ 67%) and underwent mitotic slippage, indicated by the tetraploid state, loss of mitotic markers cyclin B1, phosphorylated histone H3 Ser10, and spindle assembly checkpoint activity (BubR1) **(Supplementary Figure S1B-D**). These post-slippage cells were confirmed to have entered senescence, indicated by the discernible increase in senescence-associated beta-galactosidase (SA-βgal) activity, an enlarged flattened cell morphology, decreased Lamin B1 expression, hypo- phosphorylation of retinoblastoma protein (Rb-Ser780), and increased expression of p21, phosphorylated-p53, and DNA damage marker γ-H2AX **(Supplementary Figure S1E, F**). Persistence of senescence-associated markers in cells cultured in the absence of drug for a further three days (D6), confirmed a bona fide TIS. Similar results were also observed for CAL-51 cells **(Supplementary Figure S1G-L)**.

To test whether PTX-induced senescent MDA-MB-231 cells could influence tumour growth in xenograft mice, cycling-competent (Cyc) MDA-MB-231 H2B-mCherry cells were subcutaneously injected into immunodeficient NOD/SCID mice either alone or admixed with MDA-MB-231 H2B- GFP cells rendered senescent by prior treatment with PTX (TIS cells) **(Fig. 1A).** As shown in **Fig. 1B**, co-administration of TIS cells significantly accelerated tumour growth, compared to Cyc cells alone, indicating that TIS MDA-MB-231 cells were capable of promoting tumourigenic potential of proliferating cells *in vivo*. This was not due to senescence bypass or escape as the majority of cells dissociated from these tumours expressed H2B-mCherry **(Fig. 1C)**. Importantly, the observation that TIS cells alone did not exhibit tumour growth **(Fig. 1B)** suggested that the accelerated tumour growth was likely attributable to a pro-tumourigenic microenvironment engendered by the co-injected cells. This prompted us to analyse possible changes in the expression of secreted soluble mediators (sSASP) derived from TIS cells since pro-tumourigenic paracrine effects of TIS cells are known to be mediated via their secretome (Bavik *et al*., 2006; Faget *et al*., 2019; Jakhar *et al*., 2018). To this end, Luminex- based analysis of conditioned media (CM) derived from TIS MDA-MB-231 cells (TIS-CM) compared with DMSO-treated cells (Control-CM) at Day 6 **(Fig. 1D schematic)** revealed increased secretion of cytokines and chemokines endowed with pro-tumourigenic properties, such as IL-1α, IL1β, IP-10 (CXCL10), MCP-1 (CCL2), IL-4, GROα (CXCL1), RANTES (CCL5), and platelet derived growth factor AA (PDGF-AA) and PDGF-BB **(Fig. 1E)**.

**Figure 1.**
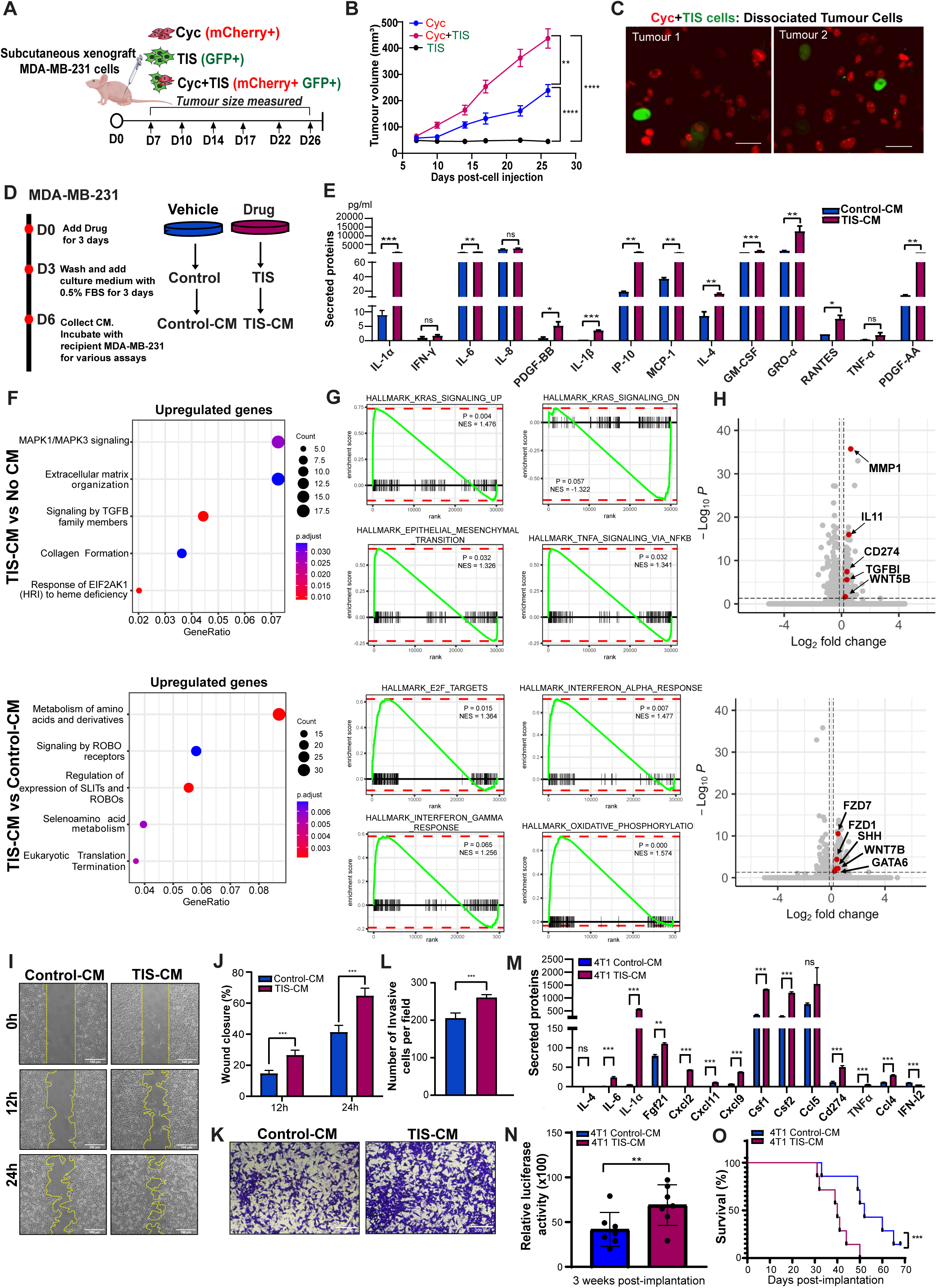
Paclitaxel-induced senescent secretome accelerates tumourigenesis in TNBC cells. (**A**) Schematic diagram of tumour growth assessment by subcutaneous injection into flank region of NOD/SCID mice of (i) non-senescent cycling-competent (Cyc) MDA-MB-231-mCherry+ cells (ii) cyc mCherry+ cells and TIS GFP+ cells (ii) TIS GFP+ cells alone. (**B**) Tumour volume (mm³) was measured at indicated timing (n = 8-10 mice per group). (**C**) Representative images for dissociated tumour cells with cyc mCherry+ and TIS GFP+ cells. (**D**) Schematic diagram of TIS induction for conditioned media (CM) collection from synchronised MDA-MB-231 cells treated with 100 ng/ml of nocodazole (Noc) or DMSO vehicle control; D:Day. (**E**) Luminex cytokine analysis at Day 6 expressed in pg/ml concentration. (**F**) REACTOME enrichment analysis of differentially expressed genes (DEGs) from MDA-MB-231 cells pre-conditioned with TIS-CM vs No CM (top) and TIS-CM vs Control-CM (bottom) depicting top five upregulated pathways. (**G**) Gene set enrichment analysis (GSEA) of expression datasets for cells as per (F) using hallmark gene sets obtained from Molecular Signature Database (MSigDB). (**H**) (top) Volcano plot of DEGs in cells incubated with TIS-CM vs No-CM, vertical dotted line denotes pLog_2_FC ≤ -0.15; Log_2_FC ≥ 0.15, horizontal dotted line denotes Padj ≤ 0.05. (bottom) Volcano plot of DEGs for cells incubated with TIS-CM vs Control-CM, vertical dotted line denotes Log_2_FC ≤ -0.15; Log_2_FC ≥ 0.15, horizontal dotted line denotes Padj ≤ 0.05. (**I**) Representative images and **(J)** quantification of wound closure after incubation with Control-CM or TIS-CM. Scale bar: 100 µm. (**K**) Representative images and **(L)** quantification of pre-conditioned cells that invaded transwell after 72h incubation per field.Scale bar 200 µm. **(M)** Cytokine analysis of CM collected from paclitaxel-treated 4T1 cells (TIS-CM) and DMSO-treated 4T1 cells (Control-CM) expressed in pg/ml concentration. **(N)** Quantitation of relative luciferase activity by bioluminescence imaging at 3 weeks post-implantation compared to 24 hours post-injection. (n=7 mice per group). **(O)** Kaplan-Meier survival curve with two-sided log-rank test conducted to determine statistical significance. Data are mean ± s.d. of 3 independent experiments. *p < 0.05, **p < 0.01, ***p < 0.001, ****p < 0.0001, and n.s. non significance by Student’s *t*-test & ANOVA.

To determine the effect of the TIS secretome on proliferating cells *in vitro*, recipient MDA-MB-231 cells were incubated with culture media alone (No CM), Control-CM or TIS-CM for 48 hours and processed for RNA-sequencing. Data analyses identified 408 genes that were differentially expressed between TIS-CM and No CM conditions, and 1064 genes between TIS-CM and Control- CM conditions **(Supplementary Table S1).** In line with the *in vivo* phenotype of accelerated growth, gene expression pattern changes of TIS CM-recipient cells were linked to aberrant growth factor signaling, metabolic pathways and pro-inflammatory-related responses consistent with a pro- tumourigenic phenotype **(Figs. 1F-1H)**. REACTOME pathway analyses of the differential expressed genes (DEGs) for TIS-CM relative to No CM revealed upregulation of genes associated with oncogenic signalling, cytokine signalling, and extracellular matrix remodelling **(Fig. 1F top)**, while that for TIS-CM relative to Control-CM were associated with upregulation in amino acid metabolism and SLIT-ROBO signalling pathways **(Fig. 1F bottom)**. In addition, Gene Set Enrichment Analyses (GSEA) using hallmark gene sets obtained from Molecular Signature Database (MSigDB) for the TIS-CM group showed significantly increased enrichment for oncogenic KRAS signalling, epithelial- to-mesenchymal transition (EMT) and cancer-promoting cytokine TNFα signalling than the No CM group **(Fig. 1G top)**, while the TIS-CM group compared to Control-CM group showed higher enrichment for targets of cell cycle regulator E2F, Type 1 Interferon response IFN-α and Type 2 Interferon response IFN-γ, and oxidative phosphorylation **(Fig. 1G bottom)**. Consistent with this, cells incubated with TIS-CM showed a significantly higher expression of genes implicated in the inflammatory response such as matrix metalloproteinase-1 (MMP1, top ranked gene), interleukin (IL)-11, programmed death-ligand 1 CD274, cytokine Transforming Growth Factor Beta 1 (TGFβ1), and the non-canonical Wnt ligand WNT5B compared to No CM **(Fig. 1H top)**. In addition, TIS-CM recipient cells showed upregulation related to growth factor signalling genes such as FGFs and EGFR, as well as angiogenic signalling genes such as FOSL1 and VEGFs compared to controls **(Supplementary Figure S2A)**. The hyperactivity of these pro-tumourigenic pathways suggested potential drug vulnerabilities such as dasatinib, digoxin and the sunitib variant SU11652 among others, as determined by our Connectivity Map (CMAP) database analysis of top perturbation hits **(Supplementary Figure S2B).** Notably, genes associated with the Wnt signalosome such as FZD1, FZD7, SHH, WNT7B and GATA6 were significantly increased in TIS-CM recipient cells compared to Control-CM **(Fig. 1H bottom)**. Wnt signalling pathway activation is known to promote cancer pathogenesis and has been reported to accompany chemotherapy-induced SASP and EMT gene expression (Basu *et al*, 2012). To assess migratory and invasive capabilities mediated by TIS-induced SASP factors in a paracrine manner, classical wound healing and transwell invasion assays were used. MDA-MB-231 cells incubated with TIS-CM promoted wound closure at a rate faster than Control- CM **(Fig. 1I, J)**, while a dramatic increase in number of cells exposed to TIS-CM compared with Control-CM invaded the bottom of the filter chamber **(Fig. 1K, L)**, confirming pro-migratory and invasive properties of the TIS secretome.

To test if the secretome of senescent murine TNBC mammary gland adenocarcinoma 4T1- 12B cells could promote tumour growth in mice, proliferating 4T1 cells expressing luciferase (4T1- Luc) were incubated for nine days with CM derived from nocodazole-treated 4T1 cells confirmed to be senescent *in vitro* **(Supplementary Figures S1M-R, Fig. 1M)**, and subsequently injected into the mammary gland of syngeneic immunocompetent BALB/C mice. As shown in **Figure 1N**, three weeks after implantation, tumours in mice injected with 4T1 cells pre-conditioned with TIS-CM exhibited a higher luciferase activity via bioluminescence imaging compared to Control-CM, suggesting greater tumour burden. Consistent with this, Kaplan-Meier survival curves for these tumour-bearing mice demonstrated that mice injected with TIS-CM-recipient cells displayed lower percentage of survival probability compared to control **(Fig. 1O)**.

### TIS-associated sEVs derived from TNBC cells impair tumour growth

Given that sEVs are entities inherent within the SASP, we postulated that sEVs derived from therapy-induced senescent TNBC cells (TIS-sEVs) might also confer tumorigenesis. This expectation was also consistent with a previous report demonstrating that EVs purified from non-transformed senescent cells could confer proliferative effects on several types of cancer cells (Takasugi *et al*., 2017).

sEVs isolated from TIS MDA-MB-231 cells using sequential ultracentrifugation (UC) followed by size-exclusion chromatography (SEC) **(Fig. 2A schematic)**, were enriched for the sEV-associated proteins TSG101, CD81, Alix and CD63, as shown by immunoblot, ELISA and flow cytometric analysis of SEC fractions 7-12 **(Fig. 2B, Supplementary Figure S3A, B)**. Nanoparticle tracking analysis (NTA) reporting a modal size of 100 and 150 nm **(Fig. 2C)**, and transmission electron microscopy (TEM) imaging **(Fig. 2D)** further confirmed these were *bona fide* sEVs. Consistent with previous reports (Lehmann *et al*, 2008; Takasugi *et al*., 2017), sEV secretion was more abundant in TIS cells compared to DMSO-treated control **(Fig. 2B, C)**.

**Figure 2.**
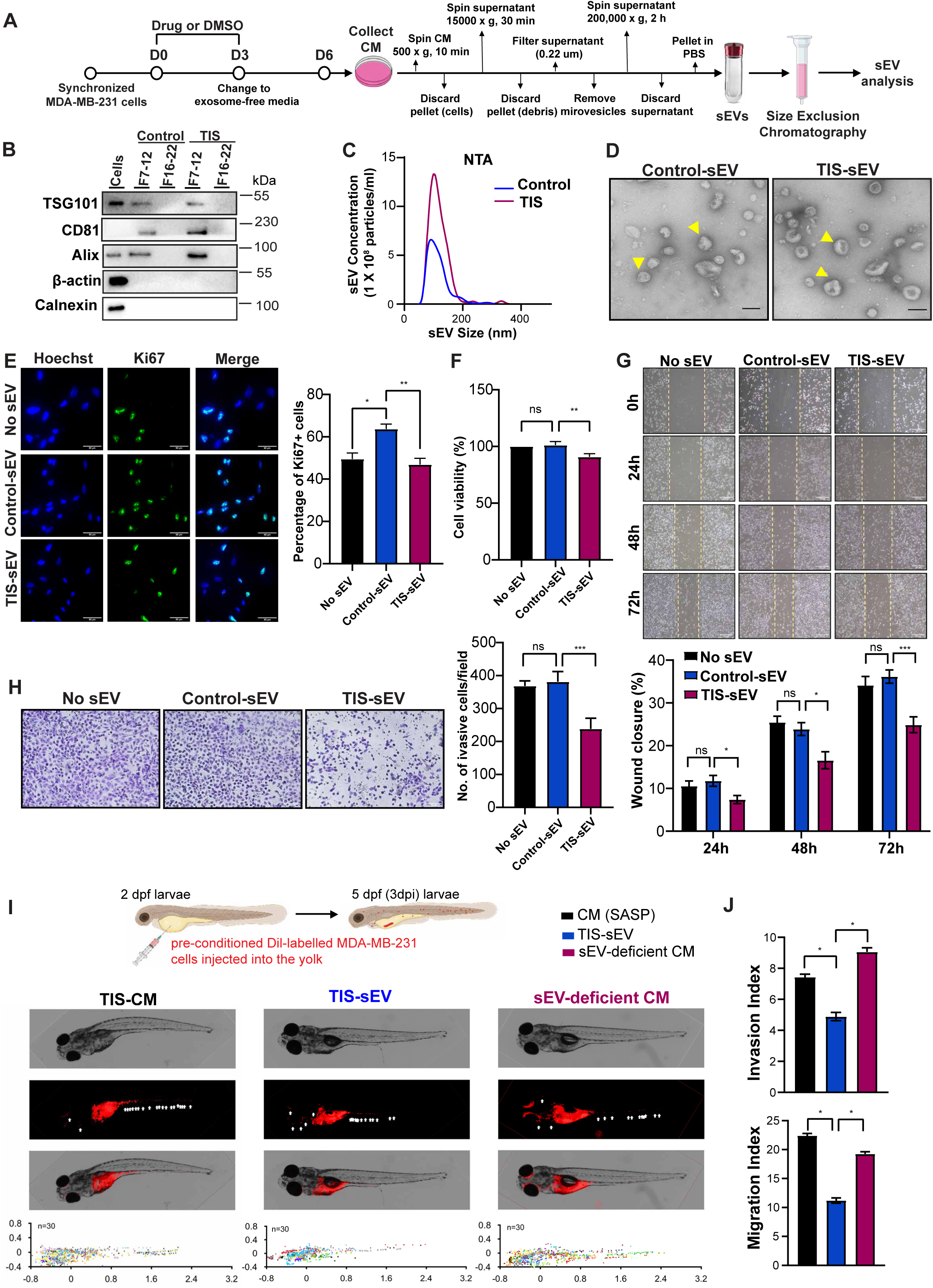
sEVs from TIS MDA-MB-231 cells attenuate invasive and migratory capacity. (**A**) Experimental design of sEV isolation from Control (DMSO-treated) and TIS (Noc-treated) MDA-MB-231 breast cancer cells using differential ultracentrifugation (UC) and size exclusion chromatography (SEC). **(B)** Immunoblot detection of eluted sEV fractions (SEC fraction 7-12), eluted protein fraction (SEC fraction 16-22), cell lysate for EV markers (Alix, CD81, TSG101) and negative controls (β-actin, calnexin). (**C**) sEV size and number were quantified using nanoparticle tracking analysis (NTA). Representative concentration vs size plot for Control-sEV and TIS-sEVs. (**D**) Representative transmission electron microscopy images of sEVs (yellow arrows) from Control and TIS cells. Scale bar: 100 µm. (**E**) Representative immunofluoresence images and quantification of Ki67-positive MDA-MB-231 cells after 72h incubation with No sEV, Cont sEV, and TIS sEV (2 x 10^9^ particles/ml). Scale bar: 50 µm. (**F**) Cell viability MDA-MB-231 cells after 72h incubation with No sEV, Control-sEV, and TIS-sEV (2 x 10^9^ particles/ml). (**G**) Representative images and quantification of wound closure in recipient MDA-MB-231 cells after 72h incubation with No sEV, Control-sEV, and TIS-sEV (2 x 10^9^ particles/ml). Scale bar: 100 µm. (**H**) Representative images and quantification of invasive recipient MDA-MB-231 cells per field after 72h incubation with No sEV, Control-sEV, and TIS-sEV (2 x 10^9^ particles/ml). Scale bar: 200 µm. (**I**) Schematic diagram of MDA-MB-231 cells flourescent-labelled with Dil (red) incubated with TIS-CM (SASP), TIS-sEV (2 x 10^10^ particles/ml) or sEV-deficient CM injected into yolk at 2 days post-fertilisation (dpf) zebrafish larvae. At 3 days post injection (dpi), zebrafish xenografts were evaluated for metastasis. Representative images of 5dpf (3dpi) larvae with tumour cells highlighted by the red fluorescence. White arrows indicate disseminated tumour foci. Tumour foci were presented as scatterplot, while ‘n’ denotes the number of injected embryos from three biological replicates. (**J**) Quantification of metastatic ability of injected MDA-MB-231 cells by Invasion index (no. of metastases formed per 100 cells injected) (left) and Migration index (distance travelled by metastases as % of maximum distance cells can migrate). Data are mean ± s.d. of 3 independent experiments. *p < 0.05, **p < 0.01, ***p < 0.001 and n.s. non significance by ANOVA.

To assess tumourigenic impact of TIS MDA-MB-231-derived sEVs (TIS-SEVs) on recipient human breast cancer cells, we first examined extent of cell viability and proliferation. To this end, recipient MDA-MB-231cells were co-cultured with TIS-sEVs and compared to uptake of Control-sEVs (sEVs from DMSO-treated cells) or culture media alone (No sEV) for three days. sEV uptake at Day 3 by MDA-MB-231 recipient cells was confirmed by their labelling with the PKH67 membrane dye **(Supplementary Figure S3C, D)**. Surprisingly (and strikingly), recipient cells with TIS-sEVs (2X10^9^ particles) showed an unexpected decrease in cell proliferation and viability compared to Control-sEVs (**Fig. 2E, F)**, indicating an anti-tumorigenic phenotype contrary to our original hypothesis. Additionally, classical wound healing and transwell assays with recipient MDA-MB-231 cells harbouring TIS-sEVs revealed a significant reduction in migratory and invasive capabilities compared to Control-sEVs **(Fig. 2G, H)**. To investigate whether this unexpected finding was also observed *in vivo*, we assessed impact of TIS-sEV on metastasis *in vivo* using the zebrafish model due to its transparency which enables visualisation of cancer cells as they disseminate, and renders cell engraftment and migration readily detectable and quantifiable. MDA-MB-231 cells were incubated with TIS-CM, TIS-sEV, or sEV-deficient CM (derived from TIS MDA-MB-231 cells treated with exosome biogenesis inhibitor GW4869) **(Supplementary Figure S3E, F)** and injected into zebrafish embryos. As shown in **Fig. 2I, J**, zebrafish harbouring TIS-sEV recipient cells exhibited lower invasion and migration compared to TIS-CM cells, indicating that TIS-sEVs conferred reduction of tumourigenic properties *in vivo.* Interestingly, removing sEVs from the SASP led to an increased invasive index, supporting the notion that the TNBC-derived TIS-sEVs represent an inherent “anti- tumourigenic” component within the “largely tumourigenic” cancer cell-derived SASP.

We wondered if the anti-tumourigenic outcome was dependent on p53 status of recipient cells. To test this, we incubated TNBC p53-wildtype Cal-51 recipient cells with TIS MDA-MB-231 cells-derived CM or sEVs **(Supplementary Figure S4A).** Both wound healing and transwell assays confirmed decreased tumourigenic potential following uptake of TIS-sEVs compared to TIS-CM, and Control-sEVs **(Supplementary Figure S4B, C),** demonstrating that alterations in p53 status of TNBCs had null impact. Interestingly, uptake of TIS-sEVs in estrogen-receptor positive (ER+) MCF-7 breast adenocarcinoma cells showed negligeable invasion and migration capabilities **(Supplementary Figure S4D-F),** suggesting that sEV-conferred impact may possibly be subtype-dependent.

To evaluate impact of murine TIS-sEVs during tumour progression *in vivo*, isolated TIS 4T1- derived sEVs **(Supplementary Figure S5)** were administered via six repeated intratumoural injections at 2-day intervals in Balb/c mice bearing 4T1 luciferase-expressing mammary tumours **(Fig. 3A)**. Consistent with phenotypes observed with human TNBC cell-derived TIS-sEVs, murine TIS-sEVs reduced tumour growth compared to Control-sEVs (sEVs derived from saline-treated cells) **(Fig. 3B-E)**. RNA sequencing analysis of tumour-derived cells harvested from mice harbouring TIS- sEVs compared to Control-sEVs and vehicle, revealed downregulation of tumourigenic-related gene expression in GO processes such as oxidative phosphorylation, oncogenic signalling, DNA repair and mitotic pathways **(Supplementary Table S2, Fig. 3F, G)**. Additionally, GSEA-based approach further confirmed anti-tumorigenicity as tumour-derived cells with TIS-sEVs displayed downregulation of mammary cancer stemness and invasiveness genes compared to Control-sEVs **(Fig. 3H)**, with the TIS-sEV vs Saline group showing downregulation of Wnt pathway and upregulation of immune response gene sets, such as T cell receptor signalling and antigen-presentation **(Fig. 3I).** Immunohistochemistry analyses comparing TIS-sEV vs Control-sEV fresh-frozen tissue sections demonstrated that cells with TIS-sEVs displayed lower proliferative potential (Ki67), and increased NF-κB-p65 activity, NKG2D ligand expression, DNA damage and p53 expression **(Fig. 3J)**. TIS-sEVs cells also showed robust senescence-associated (SA) β-galactosidase activity, suggesting tumour regression induced by murine TIS-sEVs *in vivo* is senescence-mediated.

**Figure 3.**
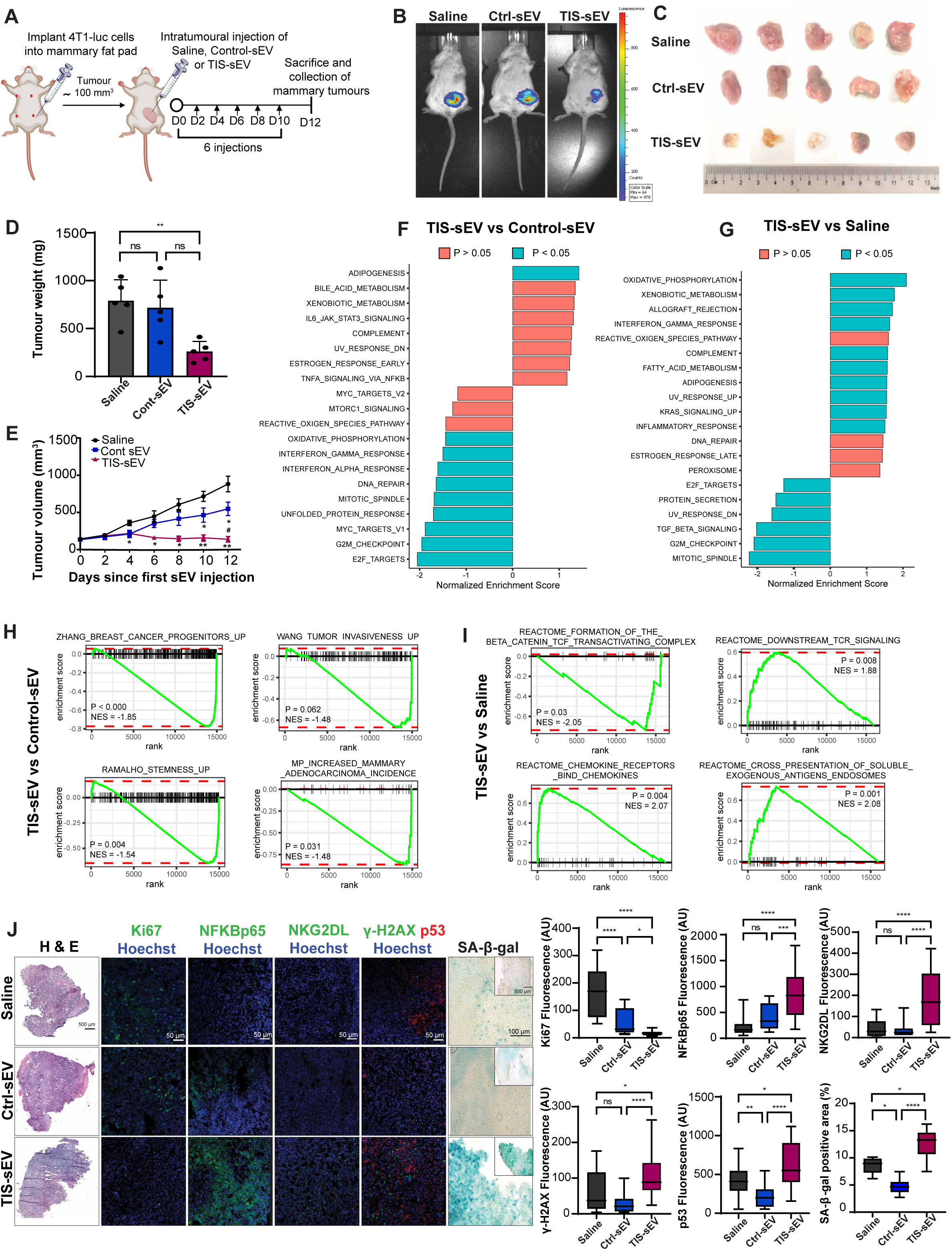
TIS derived-sEVs from 4T1 murine cancer cells reduces syngeneic tumour growth *in vivo*. **(A)** Schematic diagram of in sEV uptake in mice to assess tumour growth. BALB/c mice bearing 4T1-luc tumours in their mammary fat pads were injected intratumourally at indicated timings with 4 × 10^9^ 4T1 sEV dissolved in 50 μl PBS. n=5 mice per treatment group. **(B)** Bioluminescence imaging for mice at 12 days post-injection. **(C)** Representative images of xenograft tumours harvested from mice after 12 days of indicated treatment conditions. **(D)** Tumour weights measured at the end of the experiment. Dots represent individual tumour weights. **(E)** Tumour volumes were calculated by LxWxW/2. **(F)** HALLMARK GSEA plots of DEGs from TIS-sEV vs Control-sEV expression dataset for mice-bearing 4T1 tumour cells. **(G)** HALLMARK GSEA plots of DEGs from TIS-sEV vs Saline expression dataset for mice-bearing 4T1 tumour cells. **(H)** GSEA of TIS-sEV vs Control-sEV expression dataset using stemness and invasion-related gene sets obtained from MSigDB. **(I)** GSEA of TIS-sEV vs Control-sEV expression dataset using REACTOME gene sets. **(J)** Representative images and quantification of independent tumour sections from mice treated as per (A) showing haematoxylin and eosin (H&E), immunohistochemistry and SA-β-gal staining. Insets show tissue section from which region of interest was magnified. Data shown are mean ± SEM (n = 5 mice per group) for all experiments. *p < 0.05, **p < 0.01, ***p < 0.001, ****p < 0.0001, #p<0.05 for TIS-sEV vs Control-sEV, and n.s. non significance by ANOVA.

### TIS-associated sEVs confer genome instability and ROS-driven senescence

To better understand the biological consequences of TIS-sEV uptake, we performed bulk RNA-sequencing from recipient MDA-MB-231 cells following three-day incubation with TIS-sEV vs Control-sEVs, which revealed 74 differentially-regulated transcripts. A decrease in expression of stemness and EMT regulators was observed in TIS-EV recipient cells compared to control **(Fig. 4A, B Supplementary Table S3,)**. Notably, RNA-sequencing analysis showed downregulation of the mitochondrial superoxide dismutase 2 (SOD2) which is known to scavenge superoxide radicals and balance intracellular ROS levels, thereby preventing oxidative stress damage **(Fig. 4A, B)**. This suggested presence of high ROS levels after TIS-EV uptake. To confirm this, we measured ROS post-EV uptake by assessing abundance of SOD2 as well as the nuclear factor erythroid 2-related factor 2 (NRF2) that is stabilised in response to ROS. Immunoblotting confirmed increased ROS levels as shown by higher NRF2 levels and lower SOD2 levels in TIS-EV recipient cells compared to control **(Fig. 4C)**. This was also corroborated by flow cytometric detection using the fluorogenic dye DCFDA that detects ROS **(Fig. 4D),** as well as quantitation of reduced Glutathione levels by measuring ratio of intracellular reduced glutathione (GSH) to oxidised glutathione (GSSG) **(Fig. 4E)**, which decreases in response to ROS. This ratio decrease in TIS-sEV recipient cells could be prevented by pre-treatment of recipient cells with N-Acetyl-l-cysteine (NAC, a ROS scavenger) for 24 h prior to TIS- EV uptake **(Fig. 4E)**. We next asked if the observed ROS elevation was accompanied by concomitant DNA damage and p53 activation. Immunoblotting and immunofluorescence (IF) microscopy showed increased DNA damage (γH2AX) and p53 levels at Day 4 in TIS-sEV recipient cells compared to controls **(Fig. 4F, G)**, suggesting that TIS-EVs conferred genomic instability. To test this further, we examined frequencies of micronuclei, a marker of genomic instability (Luijten *et al*, 2018) by IF. Recipient cells bearing TIS-EVs showed a two-fold increase in micronuclei compared to control **(Fig. 4H)**. Notably, all micronuclei were found to be acentric demonstrating that chromosome fragmentation, and not whole chromosome missegregation, was elicited by sEVs. Since increased ROS can trigger senescence (Robert *et al*, 2024), we checked for senescence induction by SA-β-gal staining. As shown in **Figure 4I**, TIS-sEV recipient cells showed increased number of SA-β-gal positive senescent cells at Day 9 compared to control. This could be reversed by pre-treatment with NAC, indicating that TIS-sEVs are likely to confer ROS-mediated senescence.

**Figure 4.**
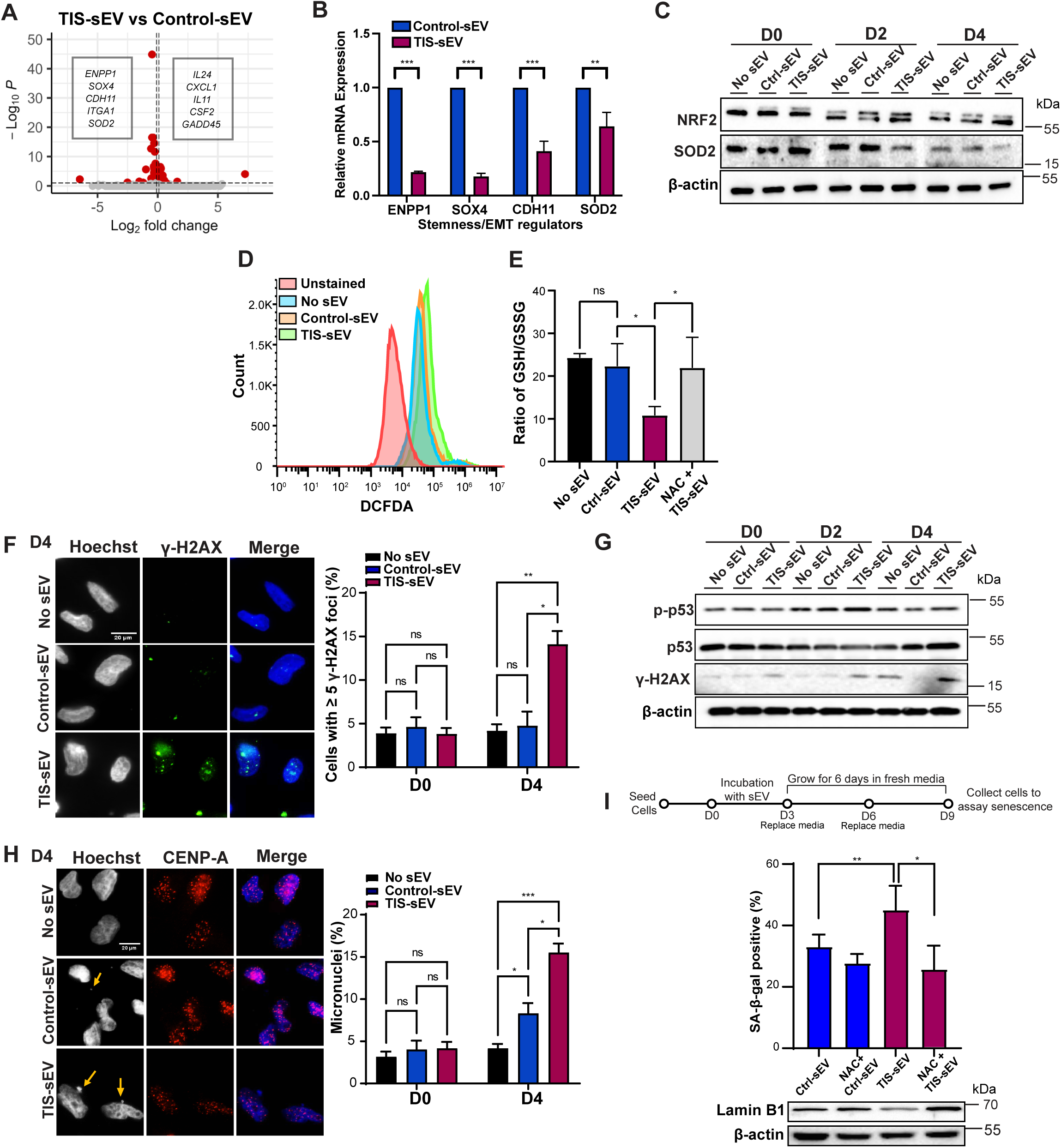
TIS-sEV confer elevated ROS, DNA damage and micronuclei formation. **(A)** Volcano plot of differentially expressed genes in recipient MDA-MB-231 cells incubated for three days with Control-sEV or TIS-sEV (2 x 10^9^ particles/ml), vertical lines denote Log2FC ≤ -0.1; Log2FC ≥ 0.1, horizontal line denotes Padj ≤ 0.1. **(B)** Validation of ENPP1, SOX4, CDH11, and SOD2 expression by qPCR in cells incubated for three days with sEVs. **(C)** Assessment of ROS-associated proteins NRF2 and SOD2, in cells incubated with No sEV, Control-sEV or TIS-sEV at Day 0 (D0; pre-incubation), Day 2 (D2), and Day 4 (D4) by Western blotting. **(D)** Flow cytometry analysis of cells stained with DCFDA two days after incubation with sEVs. **(E)** Measurement of GSH/GSSG ratio in cells pre-treated with 5mM NAC for 24 before two day-incubation with sEVs. **(F)** Representative immunofluorescence (IF) images and quantification of γ-H2AX-positive MDA-MB-231 cells incubated with sEVs at Day 4 (D4). Scale bar: 20 µm. **(G)** Assessment of DNA damage-associated proteins phosphorylated p53, p53, and γ-H2AX in cells incubated with sEVs at Day 0 (D0; pre-incubation), Day 2 (D2), and Day 4 (D4) by Western blot. **(H)** IF staining of centromere protein A (CENP-A) and quantification of acentric micronuclei in MDA-MB-231 cells incubated with sEVs at Day 4 (D4). Scale bar: 20 µm. (**I**) Quantification of SA-β-gal positive cells at Day 9 (schematic) following incubation with No sEV, Control-sEV or TIS-sEV. Cells were pre-treated with 5mM NAC for 24 prior to three-day incubation with sEVs. Western blot shows detection for Lamin B1 levels. All experiments were performed in triplicates. *p < 0.05, **p < 0.01, ***p < 0.001, and n.s. non significance by ANOVA.

### sEV-associated DKK1 suppresses tumourigenic phenotypes and promotes cytokine production via NFkB-p65/SOCS3 signaling

To identify the protein repertoire within TIS-sEVs that contribute to the anti-tumorigenic paracrine impact, a comparative proteomics approach was used to assess sEV proteins isolated from control and TIS MDA-MB-231 cells at Day 6 (as per **Fig. 2A**). Using tandem mass tag labelling-based mass spectrometry, we identified 152 significant differentially abundant proteins between TIS-sEVs and Control-sEVs, with 69 downregulated and 83 upregulated proteins **(Fig. 5A, Supplementary Table 4)**. We compared proteins detected in our proteomics analysis with the protein cargo in the Exocarta database **(Supplementary Figure S6)**. 82 TIS-sEV proteins from our experiments were common with the top 100 proteins reported on ExoCarta, GO CC and GO MF overrepresentation analyses pointed to a predominantly vesicular origin and extracellular receptor-like characteristic to be overrepresented in upregulated and downregulated proteins, as evidenced by ‘secretory granule lumen’, ‘cytoplasmic vesicle lumen’, ‘vesicle lumen’, ‘cadherin binding’, and ‘extracellular matrix binding’ being enriched in both groups **(Supplementary Figure S6 C, D)**. In addition, we also found ‘MHC class II protein complex binding’ to be enriched in the upregulated group, while hits that pertained to presence of DNA, such as ‘chromatin DNA binding, ‘nucleosome’, and ‘DNA packaging complex’ to be enriched in the downregulated group, highlighting various entities to be present within the sEVs **(Supplementary Figure S6 C, D)**.

**Figure 5.**
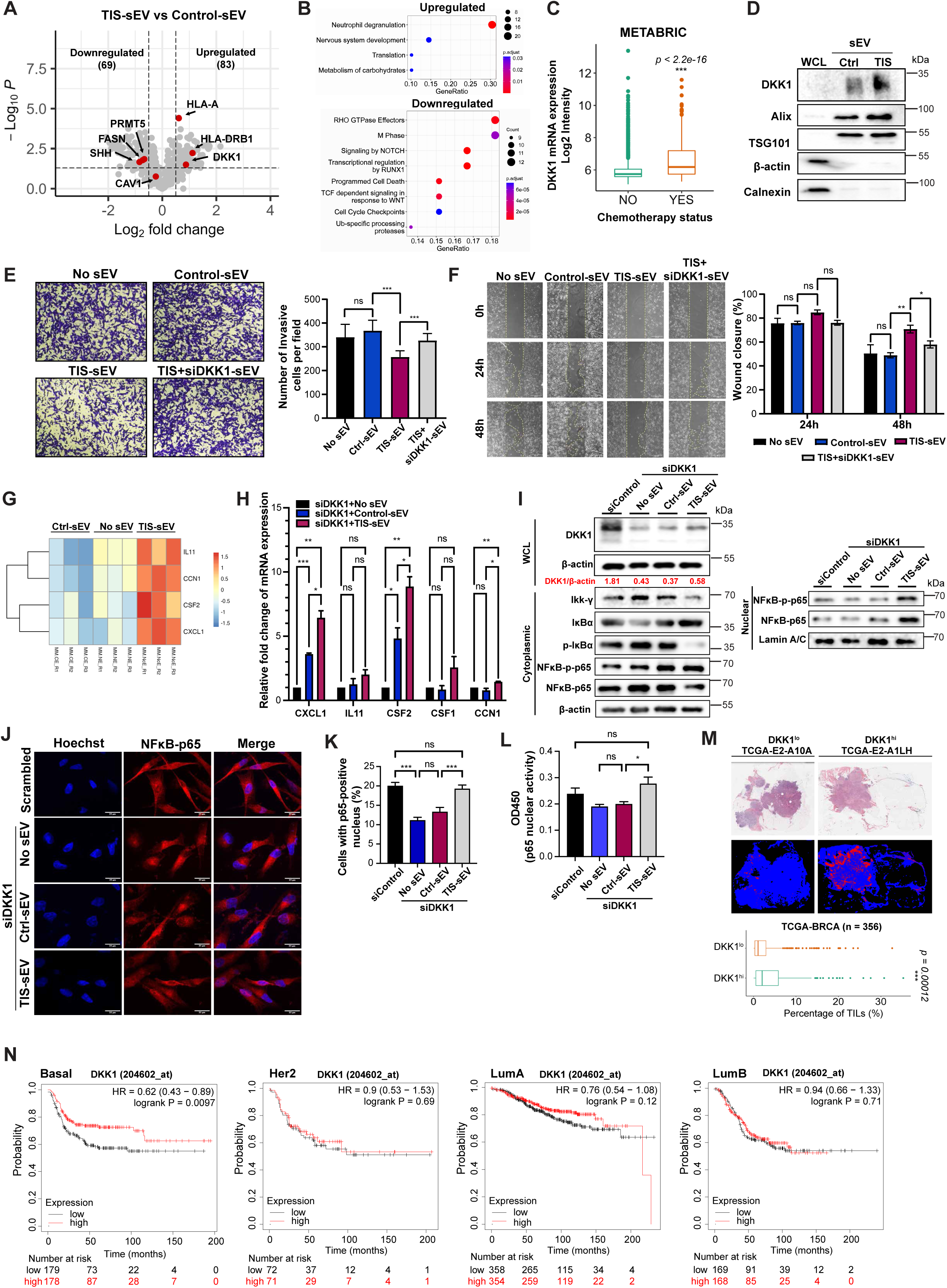
TIS sEV-enriched DKK1 increases p65 activity and inflammatory cytokine production. (A) Volcano plot of differentially abundant proteins in TIS vs Control-sEV, vertical lines denote Log2FC ≤ -0.5; Log2FC ≥ 0.5, horizontal line denotes Padj ≤ 0.05. **(B)** REACTOME enrichment analysis of differentially abundant proteins identified in TIS vs Control-sEV. **(C)** DKK1 mRNA expression levels across chemotherapy status in METABRIC cohort. **(D)** Immunoblotting of whole cell lysates (WCL) and sEVs for DKK1, β-actin, Calnexin, and sEV markers (Alix and TSG101). **(E)** Representative images and quantification of recipient cells that invaded transwell after 72h incubation with No sEV, Control-sEV, TIS-sEV, or TIS+siDKK1-sEV (2 μg/ml). **(F)** Representative images and quantification of wound closure of recipient MDA-MB-231 cell after incubation with No sEV, Control-sEV, TIS-sEV, and TIS+siDKK1-sEV (2 μg/ml). **(G)** Heatmap of significant differentially expressed cytokines, as identified by RNA-seq, between cells incubated with No sEV, Control-sEV, and TIS-sEV (2 x 10^9^ particles/ml) for 72 hours. **(H)** RT-PCR analysis of mRNA expression levels of CSF1, CSF2, CCN1, CXCL1, and IL11 in siControl (scrambled) vs siDKK1 cells. **(I)** Immunobloting of DKK1 in whole cell lysate and NFκB pathway associated proteins in cytoplasmic and nuclear fraction of siControl cells and sEV cells with siDKK1 knockeddown background after 72h incubation with sEVs (2 μg/ml). **(J)** Representative images of p65 immunofluorescence in siControl cells and cells with DKK1 knockdown background after 72h incubation with sEVs (2 μg/ml). **(K)** Quantification of positive nuclear p65 staining in siControl cells and cells with DKK1 knockdown background after 72h incubation with sEVS (2 μg/ml). **(L)** Analysis of p65 activity in siControl cells and cells with DKK1 knockdown background after 72h incubation with sEVs (2 μg/ml) via colorimetric transcription factor assay kit. **(M)** Representative H&E staining and computational TIL staining images of breast cancer tumours from TCGA-BRCA, retrieved from the CANCER Digital Slide Archive and TCIA respectively, quantification of estimated TIL proportions in DKK1^hi^ (top 25th percentile) and DKK1^lo^ (bottom 25th percentile) breast cancer tumours are shown below representative images. **(N)** Kaplan-Meier survival curve from pooled GEO database obtained via *KMplotter* indicating relapse-free survival (RFS) of breast cancer patients with DKK1-high (top 25th percentile) vs DKK1-low (bottom 25th percentile) across subtypes as identified by PAM50 and St. Gallen criteria. *p < 0.05, **p < 0.01, ***p < 0.001, ****p < 0.0001 and n.s. non significance by Student’s t-test & ANOVA.

Interestingly, amongst the upregulated hits, we found the major histocompatibility complexes (MHC), HLA-A and HLA-DRB1, and the secreted Wnt/β-catenin signalling antagonist, **Dickkopf-1 (DKK1)**, to be highly-enriched in the TIS-sEVs. In line with the latter, downregulated hits included the protein arginine methyltransferase PRMT5, known to activate WNT/ β-catenin signalling in breast cancer cells via epigenetic silencing of DKK1 and DKK3 (Shailesh *et al*, 2021). REACTOME enrichment analysis also identified the downregulation of other signaling proteins pathways such as those related to RHO GTPase, NOTCH, RUNX1 and WNT pathways in TIS-sEVs **(Fig. 5B)**. To identify a clinically-relevant pathway by which TIS-sEVs might exert their tumour-suppressive effect, we analysed the METABRIC breast cancer datasets which revealed a significantly higher level of DKK1 mRNA expression in patients that received chemotherapy, compared to those who did not **(Fig. 5C)**. Consistent with this, immunoblotting confirmed the higher abundance of DKK1-specific bands (made up of different protein isoforms) in TIS-sEVs established to elicit tumor suppression compared to control **(Fig. 5D).** To ascertain whether sEV-associated DKK1 contributed to the tumour-suppressive phenotype observed, TIS-sEVs were isolated from cells where DKK1 expression was “knocked down” by means of small interfering RNAs (siRNAi; DKK1-KD). As shown in **Figure 5E and 5F**, TIS-sEVs derived from DKK1-KD cells failed to hamper migratory and invasive capacity normally observed with TIS-sEVs in recipient cells, strongly suggesting that sEV-associated DKK1 played a critical role in the paracrine anti-tumourigenic outcome **(Fig. 5E, F)**.

A recent study pointed to a cell-autonomous function for DKK1 in promoting inflammatory cytokines for mounting an immune response in cancer cells (Jaschke et al, 2022). Our RNA-sequencing analysis of recipient cells following uptake of TIS-sEVs had shown increased expression of several pro-inflammatory cytokines, namely CXCL1, CSF2, CCN1 and IL11, compared to Control-sEVs and no sEV **(Fig. 5G, Supplementary Figure S7A, Supplementary Table S3)**. Interestingly, we found a positive correlation of DKK1 expression with CXCL1, CSF2, CCN1, and IL11 expression in TCGA-BRCA cohort **(Supplementary Figure S7B)**. Quantitative PCR analysis indeed confirmed that suppression of DKK1 expression curtailed the expression of CXCL1, CSF2, and CCN1, but not CSF1 and IL11, suggesting control of cytokine production via DKK1 **(Supplementary Figure S7C)**. To ascertain a possible link between DKK1 and cytokine production in TIS-sEVs, we devised a “DKK1 rescue” experiment where we depleted DKK1 in recipient cells via siRNA before co-culture with sEVs, such that supplementation of DKK1 was via sEV uptake alone. Consequently, we identified an increase in CXCL1, CSF2, and CCN1 cytokine production (but not CSF1 and IL11) in TIS-sEV recipient cells compared to Cont-sEV in DKK1 KD cells **(Fig. 5H)**. Immunoblotting confirmed increased DKK1 protein abundance in whole cell lysate, which was highest following TIS-sEV incubation in DKK1-KD cells **(Fig. 5I)**. Interestingly, recipient TIS-sEV cells exhibited increased NF-kB p65 activity in the nucleus compared to Cont-sEV in DKK1-KD cells, shown by p-p65 and p65 accumulation in nuclear fractions via immunoblotting **(Fig. 5I)**, increased nuclear colocalization of p65 via IF **(Fig. 5J, K)**, and by colorimetric p65 activity **(Fig. 5L)**. Of note, we found reduction of Ikk-γ and persistent presence of IκBα despite accumulation of nuclear p-p65 and p65 in TIS sEV recipient cells, suggesting a non-canonical p65 activation **(Fig. 5I)**.

We then sought to investigate the potential clinical impact of DKK1 by leveraging publicly available databases. To this end, we first analyzed differentially expressed genes in DKK1^hi^ (top 25th percentile) vs DKK1^lo^ (bottom 25th percentile) basal-only tumors from the TCGA-BRCA database. We found high DKK1 to be correlated with inflammatory and immune activation related pathways, as evidenced by enriched GO BP, CC, and MF terms such as ‘inflammatory response’, ‘immune response’, ‘chemotaxis’, ‘extracellular region’, ‘chemokine activity’, and ‘cytokine activity’ being upregulated in DKK1^hi^ tumors **(Supplementary Figure S7F)**. In contrast, we found enrichment of ion transport-related pathways, typically linked to tumor progression (Djamgoz *et al*, 2014), being downregulated in the DKK1^hi^ tumors **(Supplementary Figure S7F)**. To further investigate the relationship of DKK1 and tumor immune profiles, we leveraged on a convolutional neural network– based atlas by The Cancer Image Archive (TCIA) (Clark *et al*, 2013; Saltz *et al*, 2018) where we found a positive association of tumor infiltrating lymphocytes (TILs) and high DKK1 expression in human breast tumors **(Fig. 5M)**. We also leverage on publicly available single-cell atlas of human breast tumors, and interestingly found that DKK1 is primarily expressed only in breast cancer cells, specifically of Basal and Her2 subtypes **(Supplementary Figure S7E)**. This is consistent with DKK1 expression analyses using TCGA BRCA and METABRIC datasets, that also revealed significantly higher DKK1 expression in basal and Her2 subtypes, compared to Luminal A and Luminal B subtypes **(Supplementary Figure S7D)**. Notably, a meta-analysis of multiple GEO datasets, retrieved from KMplotter, showed that high expression of DKK1 may predict better relapse-free survival of patients with basal breast tumors, as identified via PAM 50 and St Gallen criteria (Coates *et al*, 2015; Parker *et al*, 2009) compared to other subtypes **(Fig. 5N)**. Taken together, our data support a role for TIS sEVs-associated DKK1 in modulating pro-inflammatory and immunogenic cytokines, resulting in anti-tumor immunity, specifically in the basal phenotype of breast cancers.

## METHODS

### Cell culture and construction of stable cell lines

Human cell lines MDA-MB-231, CAL-51, MCF- 7, RPE-1, MCF10A, and mouse 4T1 cells were purchased from American Type Culture Collection (ATCC, USA). Cells were cultured in DMEM medium (Gibco, USA) supplemented with 10% fetal bovine serum (FBS; Cytiva), 1% penicillin/streptomycin (P/S; Invitrogen) in a humidified 5% CO_2_ incubator at 37°C. The pEGFPC2-CD63 and mCherry-CD81 plasmids (Addgene, USA) were transfected into cells using Lipofectamine 2000 reagent (Invitrogen, USA) according to manufacturer’s instructions. G418 (Geneticin, Sigma-Aldrich, USA) was added 48 h post-transfection. After selection for 14 days, stable GFP-CD63 and mCherry-CD81 MDA-MB-231 cell lines were harvested for use. We have confirmed the absence of mycoplasma contamination in our cells.

### Therapy-induced senescence induction

Cells were first synchronised by serum starvation-single thymidine method prior to nocodazole or paclitaxel addition. Briefly, cells at 30%-40% confluency were grown in serum-free media for 24 h to reach G0/G1 arrest, washed with DMEM without supplements, and grown in complete media containing 3 mM thymidine (Sigma-Aldrich, USA) for 24 h to reach G1/S arrest. To resume cell proliferation, cells were washed three times with FBS/P/S-free media and grown for further 3h in normal media. Cells were then treated with antimitotic drug Nocodazole (Noc, 100 ng/ml; Sigma-Aldrich, USA) or Paclitaxel (PTX, 150 nM; Sigma-Aldrich, USA) for 72 h to achieve therapy-induced senescence (TIS) state.

### Viability assay

Cell viability and proliferation was assayed using CellTiter 96 Aqueous One Solution Reagent (Promega, USA) according to manufacturer’s instructions.

### Senescence-associated β-galactosidase (SA-β-gal) staining

SA-β-Gal activity was measured using the SA-β-gal staining kit (Cell Signaling Technology, USA) following manufacturer’s protocol. Number of blue cells (senescent) relative to the total cell number was counted in two to four different optic fields using EVOS microscope 40X objective lens (Thermo Fisher Scientific, USA). At least 200 cells were counted per sample.

### Total RNA extraction, cDNA synthesis and qRT-PCR

Total RNA was isolated using RNeasy Plus kit (Qiagen, USA). RNA quality and concentration were determined using NanoDrop 2000 system (Thermo Fisher Scientific, USA). Reverse transcription of RNA to cDNA was performed using the iScript cDNA synthesis kit (Bio-rad Laboratories, USA). cDNA was mixed with SYBR Select Master Mix for CFX (Applied Biosystems, USA) and relevant primers (Integrated DNA Technologies, USA). Quantitative real-time PCR (qRT-PCR) reaction was carried out with Step-One Plus system (Thermo Fisher Scientfic, USA).

### Flow cytometry

Cells were fixed overnight with ice-cold 70% ethanol (Sigma-Aldrich, USA). For phospho-histone-H3/propidium iodide analysis, fixed cells were washed twice in PBS with 0.5% BSA prior to incubation with 1:500 dilution of phospho-histone H3 antibody conjugated with Alexa Fluor 488 (Cell Signaling Techonology, USA) for 1 h in the dark at room temperature. The samples were then washed in 0.5% BSA in PBS and incubated with 250 µg/ml RNase A (Sigma-Aldrich, USA) and 10 µg/ml propidium iodide (PI, Sigma-Aldrich, USA) at 37°C for 30 min before analysis. FACS analyses were performed with BD Accuri C6 Flow cytometer (BD Biosciences, USA) using two-parameter scanning; FL1-H (FITC) versus FL2-A (PI) and data analysed using FlowJo software (BD Biosciences, USA).

### Immunoblotting

For total protein isolation, cells were lysed in RIPA buffer (Pierce, USA) containing protease/phosphatase inhibitor cocktail (Thermo Fisher Scientific, USA). Cytoplasmic and nuclear portion of protein were extracted by NE-PER nuclear and cytoplasmic extraction reagents (Thermo Fisher Scientific, USA). Protein concentration was measured using the Bradford assay (Bio-Rad Laboratories). An appropriate volume of Laemmli sample buffer was added to equal amounts of total protein followed by boiling for 5 min. Samples were subjected to SDS–PAGE electrophoresis before transfer to nitrocellulose membrane (Bio-Rad Laboratories, USA). After blocking for 1 h in 5% skim milk/PBST, cells were incubated overnight at 4°C in primary antibody diluted in 3% skim milk/PBST. Membranes were then washed three times with PBST and incubated with HRP-conjugated secondary antibody diluted in 3% skim milk/PBST. Signal was detected using enhanced chemiluminescence (ECL) (GE Healthcare Life Sciences, USA) and ChemiDoc imaging system (Bio-Rad Laboratories, USA).

#### Immunofluorescence

Cells grown on coverslips were fixed in PBS/4% paraformaldehyde (PFA) for 10 min at room temperature. Cells were then washed twice with PBS, permeabilised with PBS containing 0.1% Triton X-100 (PBST) and incubated in blocking buffer (PBST/10% FBS/1% BSA) for 30 min. Cells were then incubated in blocking buffer containing primary antibody overnight at 4°C. Following three washes with PBS containing 0.1% Triton X-100 (PBST), cells were incubated in blocking buffer containing secondary antibody for 2 h at room temperature. Nuclei were stained with Hoechst stain for 10 min at room temperature. Images were captured using Nikon Inverted Microscope Eclipse Ti-E (Nikon) and Orca ER CCD camera (Hamamatsu Photonics).

### Conditioned media preparation

72h after drug addition, TIS MDA-MB-231 cells were grown in reduced serum media DMEM/0.5% FBS/1% P/S for further three days. Cell culture medium was collected and centrifuged at 5000×g and the supernatant filtered with 0.22_µm pore filter (Pall Corporation, USA). This was mixed with media containing 40% FBS in a proportion of 3:1 to generate conditioned media (CM) containing 10% FBS.

### Chemokine and cytokine analyses

CM was collected from TIS MDA-MB-231 cells for chemokine and cytokine analyses using a Milliplex MAP human cytokine/chemokine magnetic bead panel (Merck Millipore) according to the manufacturer’s protocol.

### sEV isolation from conditioned media

For sEV isolation using ultracentrifugation protocol, CM was prepared after drug or DMSO treatment by incubating cells in growth media containing 5% exosome-depleted FBS (Thermo Fisher Scientific, USA) for 72 h. CM was then collected and centrifuged at 500×g for 10 min, 2,000×g for 30 min, and 15,000×g for 30 min at 4°C to remove cells and large debris. The supernatant was then filtered using 0.22 μm pore filter (Pall Corporation, USA), and EVs pelleted at 120,000×g for 2 h at 4°C. The EV pellet was washed with PBS and pelleted again by centrifugation at 100,000×g for 1 h at 4°C. Isolated EVs were further dissolved in 0.5 ml of PBS and purified using SEC-qEV2/35 nm columns (Izon Science, New Zealand) for size-exclusion chromatography (SEC). After rinsing the qEV columns with PBS, 0.5 ml of EV sample were applied on top of the column, equilibrated, and eluted with PBS in 30 fractions (0.5 ml/fractions). Two sets of four EV-rich fractions (7–12 and 16-22) were pooled and concentrated using Amicon Ultra-4 10 kDa centrifugal filter device (Merck Millipore, USA). Alternatively, sEV isolation was performed using the Total Exosome Isolation Reagent (from cell culture media) kit (Invitrogen, USA) according to manufacturer’s protocol. Briefly, CM was incubated with kit reagents at 4°C, and precipitated sEVs recovered by centrifugation at 10,000xg for 60 min.

### sEV detection and quantification

For Nanoparticle Tracking Analysis (NTA), sEVs were quantified based on size distribution using NanoSight NS300 (Malvern Instruments, UK). For data acquisition, approximately 10 µl of EV suspension diluted in final volume of 1 ml, was loaded into the sample chamber of the NanoSight using a sterile syringe. Samples were captured at 20-25°C with camera level of 13 and gain of 250–300 over 60 s in triplicates. For data analysis, a detection threshold of 3 and Blur/track length/expected size set at Auto were used. Data were analysed with the Nanosight NTA software version 3.1 (Malvern Panalytical, UK). Alternatively, EVs were quantified with the ExoELISA Complete Kit (CD63 Detection, System Biosciences, USA) according to manufacturer’s instruction. For detection and quantitation of surface CD63 by flow cytometry, SEC fractions were incubated overnight with anti-CD63 magnetic beads (Thermo Fischer Scientific, USA) with continuous rotation at 4_°C. This was followed by flow cytometric detection after staining of bead-bound sEVs with fluorescent anti-CD63 phycoerythrin (PE) (Sigma-Aldrich, USA).

### Transmission electron microscopy

Aliquots from pooled EV-rich SEC fractions (F7-12) were deposited onto formvar-coated 400 mesh copper grids for 7 min at room temperature and thereafter stained with 2% uranyl acetate (Electron Microscopy Services, USA). Grids were imaged using transmission electron microscope Tecnai T12 (Thermo Fisher Scientific, USA).

### Visualisation of EV uptake

MDA-MB-231-derived sEVs were labelled using the PKH67 Green Fluorescent Cell Linker kit (for general cell membrane labelling, Sigma-Aldrich, USA) according to manufacturer’s protocol. Cells were also stably transfected with GFP-CD63 and mCherry-CD81 plasmids (Addgene, USA) for exosomal tracking. Recipient cells were incubated with either PKH67 dye-labelled or fluorescent-labelled sEVs for 2-3 days in media supplemented with 5% exosome- depleted FBS (Gibco, USA). Cells were imaged with time-lapse microscopy or washed with PBS, fixed in 4% PFA for 15 min and examined under Nikon Eclipse Ti microscope for immunofluorescence.

### Tandem mass tag labelling and data analysis

Protein was isolated from purified EVs using Total Exosome RNA & Protein Isolation Kit (Invitrogen, Thermo Fisher Scientific) following manufacturer’s instructions. 100 µg protein of the EV lysate in 8 M urea was alkylated with dithiothreitol (DTT) and iodoacetamide (IAA), diluted with seven volumes of PBS and digested with Trypsin/Lys-C for proteolytic digestion. The reaction was quenched by heating at 60 °C. Digested proteins were desalted, dried and dissolved in 200 mM triethylammonium bicarbonate buffer (pH, 8.5). Tandem Mass Tagging (TMT) labelling was performed using Tandem Mass Tag Reagents kit (Thermo Fisher Scientific, USA) following manufacturer’s protocol. Different TMT labels were used to label different groups: TMT-126, 127, and 128 were used for cont-sEV group; TMT-129, 130, and 131 for TIS-sEV group. After labelling, samples were pooled, dried and dissolved in 0.1% trifluoroacetic acid (TFA). The samples were then desalted, dried and dissolved in 100 µL of 0.1% TFA. ExoCarta (web-based compendium of exosomal cargo) were used to identify exosomal protein cargo, with the top 100 sEV proteins then further analysed using DAVID software. Proteins identified were also subjected to pathway enrichment analyses and visualization using R v4.1.2. Gene Ontology (GO) enrichment analysis was performed by clusterProfiler v4.2.2, while Reactome enrichment analysis was by ReactomePA v1.38.0. For visualization, enrichment analyses and volcano plot were constructed using enrichplot v3.15 and EnhancedVolcano v1.14.0 respectively.

### RNA sequencing analyses

Isolated RNA samples were sent to NovogeneAIT Genomics Pte Ltd (Singapore) for sequencing. Briefly, raw reads of fastq format were firstly processed through fastp software to obtain clean reads by removing reads containing adapter, poly-N and low-quality reads from raw reads. RNA-seq reads were then aligned to the human genome using Hisat2 v2.0.5, and featureCounts v1.5.0-p3 was used to count reads numbers mapped to each gene. Differential expression analysis was performed using DESeq2. The resulting p-values were adjusted using the Benjamini and Hochberg’s approach for controlling the false discovery rate. Genes with an adjusted p-value (padj) ≤ 0.05 found by DESeq2 were assigned as differentially expressed unless otherwise stated. Resulting differentially expressed gene list was used for downstream enrichment analyses. Gene Ontology (GO) enrichment analysis was performed by clusterProfiler v4.2.2, while Reactome enrichment analysis by ReactomePA v1.38.0. For enrichment visualization, enrichment analyses and volcano plot were constructed using enrichplot v3.15 and EnhancedVolcano v1.14.0 respectively. Lastly, gene set enrichment analysis (GSEA) was performed using fgsea v4.2.

### Cellular reactive oxygen species (ROS) measurement

Cells incubated with sEV for 72h were trypsinised and washed twice with PBS. After washing, cells were stained with 20_μM of 2’,7’– dichlorofluorescin diacetate (DCFDA) (ab113851) (Abcam, USA) for 30_min at 37_°C and analysed on an Accuri C6 flow cytometer (BD Biosciences, USA) under FL-1 channel. Data were analysed using FlowJo software.

### GSH/GSSG ratio measurement

GSH/GSSG ratios were measured using GSH/GSSG-Glo™ Assay (Promega) according to the manufacturer’s protocol. Briefly, cells were lysed using Total Glutathione Lysis Reagent and Oxidized Glutathione Lysis Reagent. After lysis, Luciferin Generation Reagent was added unto the wells and plate was incubated for 30 minutes at room temperature. Subsequently, Luciferin Detection Reagent was added unto the wells, followed 15 minutes incubation and luminescence measurement.

### Wound healing assay to assess migration

Cells were seeded in 6-well plates to reach confluency. The confluent monolayer was scratched with a 200_µl pipette tip to generate a wound and incubated for 72 h with CM or sEV-containing media (2x10^9^ particles/ml). Phase-contrast images were captured at 24 h, 48 h, and 72 h timepoints under 10X objective lens and analysed using ImageJ.

### Transwell assay to assess invasion

Cells were incubated with CM or media containing sEV (2x10^9^ particles/ml) for three days. Cells were then trypsinized, resuspended in CM or sEV-containing media, seeded onto transwell inserts (8_µm pore size) pre-coated with Matrigel (BD Biosciences, USA), with 20% FBS added to the lower chamber and further incubated for 24h. Non-invasive cells were removed from the upper surface of the inserts and invasive cells on the lower surface were fixed with PFA, stained with crystal violet and observed under 10X objective lens. The number of invaded cells in six to nine random fields were counted per condition.

### Animal work

#### Quantification of cellular migration and metastasis in zebrafish (ZgraftTM)

The Zebrafish Research Facility in Acenzia is certified via Ontario Ministry of Agriculture, Food and Rural Affairs (OMAFRA). All protocols used were monitored by the Animal Health and Welfare Branch, OMAFRA. Fish in the facility were housed and maintained using Tecniplast’s ZebTec system with a controlled day-night (14 h light/10 h dark) light cycle. Embryos obtained from natural spawnings and developmental stages are reported as hours post-fertilization (hpf) at 28.5°C. sEVs were isolated from drug-treated senescent MDA-MB-231 cells. MDA-MB-231 cells were incubated with the sEVs at a concentration of 2x10^10^ particles/ml, or with conditioned media (SASP or sEV depleted-SASP) two days before injection. For collecting conditioned media with sEV-depleted-SASP, MDA-MB-231 cells after treatment with 15 μM N-Smase inhibitor GW4869 (Sigma-Aldrich, USA) for 72h, were grown in reduced serum media DMEM/0.5% FBS/1% P/S for further three days. Conditioned media was collected and processed as mentioned before. Xenotransplanted MDA-MB-231 cells were visualised in the zebrafish embryos by red fluorescence by staining with Vybrant DiI (Life Technologies, USA). Cells from each treatment were detached at 80% confluency using Trypsin- EDTA solution (Sigma-Aldrich, USA). Cells were further incubated for 45 min at 37°C in serum-free culture media containing DiI at a concentration of 5 μl dye per million cells. Cells were then washed twice and re-suspended in serum-free media for injection. Zebrafish embryos at 48 hpf were then dechorionated when required and anesthetised with tricaine (Sigma-Aldrich, USA). Using Nanoject II (Drummond Scientific, USA) injector, approximately 100 cells were injected into the yolk of each embryo. Embryos were then incubated for 2 h at 32°C and checked for successful injection. Embryos with fluorescent cells outside the yolk sac were excluded from further experimentation and analysis. Accurately-injected embryos were further incubated at 32°C in a fresh fish tank system water for 48 h and imaged. Embryos were anesthetised on a microscope slide at 48 h post-injection and imaged using Leica M165 LC. Images were gathered, aligned to a specific orientation and analysed by ImageJ software to determine metastatic tumour foci position relative to injection site (0,0 on graph). Tumour foci observed in all tested larvae, in each treatment group (presented as n on the graph), were presented as scatter plots. All foci in one larva were represented by a single colour. For consistent quantitative comparison of differentially treated cells, Invasion Index measured the ability of cells to invade, and Migration Index measured the ability of cells to migrate. These were calculated as follows: Invasion index = Number of metastases formed per 100 cells injected. Migration index = Distance travelled by metastases as percent of the maximum distance cells can migrate within the fish.

### Mice xenograft experiments

All animal experiments were conducted in accordance with the National Advisory Committee for Laboratory Animal Research (NACLAR) Guidelines and approved by the National University of Singapore (NUS) Institutional Animal Care and Use Committee (IACUC) under Protocol No. R18-0635. BALB/c mice were purchased from InVivos, Singapore. Mice were kept on a 12-h light/dark cycle with food and water provided *ad libitum* and maintained under pathogen-free conditions in the Animal Housing Unit, Comparative Medicine Department at the National University of Singapore.

Immunocompromised (NOD/SCID) Balb/c female mice were transplanted subcutaneously into the right flank with 2.5x10^6^ untreated cycling-competent (cyc) MDA-MB-231 cells alone, 2.5x10^6^ cyc MDA-MB-231 cells and 0.5x10^6^ cells paclitaxel-induced senescent MDA-MB-231 cells, or 2.5x10^6^ cells paclitaxel-induced senescent MDA-MB-231 cells alone.

NOD/SCID female mice 6–8 weeks of age were injected subcutaneously (sc) under anesthesia with 2.5 × 106 MDA-MB-231 cells on the right flank. When tumors reached a volume of about 100 mm3, mice were randomized into two groups of five and six animals, and received a total of 7 ip injections of 100 μl saline containing 100 μg either Andes-1537 (group of 5) or ASO-C (group of 6) on days 12, 14, 16, 19, 21, 23, and 26 post-cell inoculation, in a blinded fashion. Tumor growth was monitored with a caliper and tumor volumes were calculated following the formula: tumor volume = length × width2 × 0.5236. Mice were sacrificed under anesthesia on day 27 after cell injection.

### Tumour implantation, sEV challenge and bioluminescence imaging of 4T1 tumour

Fifteen mice were randomised into three groups (saline, DMSO control sEVs, and TIS sEVs, 5 mice per group) 50 μl of sterile PBS containing 1×10^5^ 4T1-12b cells were injected into the mammary fat pad of BALB/c mice. Upon reaching a tumour size of approximately 100mm^3^, 50 μl of saline, control sEVs or TIS sEVs generated from mouse breast cancer 4T1 cells at a concentration of 4×10^9^ particles/50 μl, were injected intratumourally every other day for 6 injections. Mice were weighted prior to, during and post-injection. Tumours were localized and quantitated using bioluminescence-based Xenogen IVIS Spectrum Imaging System (PerkinElmer, USA). Briefly, mice were injected with 150 µL of VivoGlo Luciferin (150 mg/kg) (Promega, USA) intraperitoneally prior to imaging and put under isoflurane anaesthesia during imaging. Tumour volumes were measured manually using a digital calliper and volume calculated as follows: V (mm^3^) = length (mm) × width (mm) × width (mm)/2). Mice were euthanized either at the end of the study or earlier if they displayed significant weight loss, signs of distress, or palpable tumours ≥ 2.0 cm in diameter. Tumours were excised and used for downstream molecular analyses.

### Mice tumour-associated immune cell characterisation

Cells were stained with flexible viability dye eFluor 506 (myeloid panel-1:1000) or near IR (lymphoid panel-1:100) (Thermo Fisher Scientific, USA). Cells were subsequently stained with the following cocktail: CD45-AF700, CD 11b-FIC, CD86-APC, CD206-PE/Cy7, MHC Class II-APC eFlour 780, LY-6G-PE, F4/80-PB, CD11c-PE eFluor 610 (Myeloid Panel) or F4/80-PE/Cy, CD278-AF647, CD8a-BV421, CD45-BV711, CD279-PE-CF594, CD3e-BUV395, CD4-AF488, CD25-PE (Lymphoid panel). Cells were analysed on a BD LSRFortessa X-20 (BD Biosciences, USA) with data analysed using FlowJo software.

### Immunohistochemistry of mice tissues

Harvested mice tissues were stored in 4% PFA overnight followed by sequential dehydration with 15% ethanol (overnight) and 30% ethanol (overnight). Tissues were then mounted on mounting blocks made from Tissue-Tek O.C.T. (Sakura) at -20C, before being sliced by cryostat at 10 uM thickness with a cryostat (Leica Microsystems CM3050S). Slices were picked up onto cover slides and rinsed by deionized water briefly followed by haematoxylin staining for 10 min and eosin counterstain for 20 seconds. Slices were then washed with 95% ethanol and xylene before being mounted in xylene-based mounting media on the coverslips. After the mice tissues were dehydrated and sliced with cryostat as per protocol above, slices were picked up on coverslips and briefly rinsed with PBS. Slices were then blocked with blocking buffer (PBS, 5% Normal Donkey Serum, 0.3% Triton-X) at 4 °C for 1 h, and subsequently incubated with primary antibodies overnight. Slides were then washed three times with PBST at 4 °C for 10 min, and incubated with blocking buffer containing secondary antibodies (1:200) at room temperature for 2 h. This was followed by three PBST washes and DAPI counterstain. Slices were then mounted onto slides with mounting media.

### Quantification and statistical analysis

All experiments were performed independently at least three times. Statistical analyses were performed using Prism 9 software (GraphPad), and statistical information pertaining to the number of replicates and significance can be found in Figure Legends. Significance levels for comparisons between two groups were determined with two-tailed Student t-test, and for comparisons of more than two groups were determined with one-way analysis of variance (ANOVA). Data were plotted as mean ± SD.

## ACKNOWLEDGEMENTS

We are grateful to Brian Kennedy, Yeong Foong May, and Vaidehi Krishnan for support and discussions, and Clarinda Lim for administrative help.

