## Supplementary figures and images for "Tumour-derived extracellular vesicles within the therapy-induced senescent secretome distinctly suppress breast cancer via DKK1-mediated inflammatory response"

### Supplemental Figure 1

#### Supplementary Figure S1

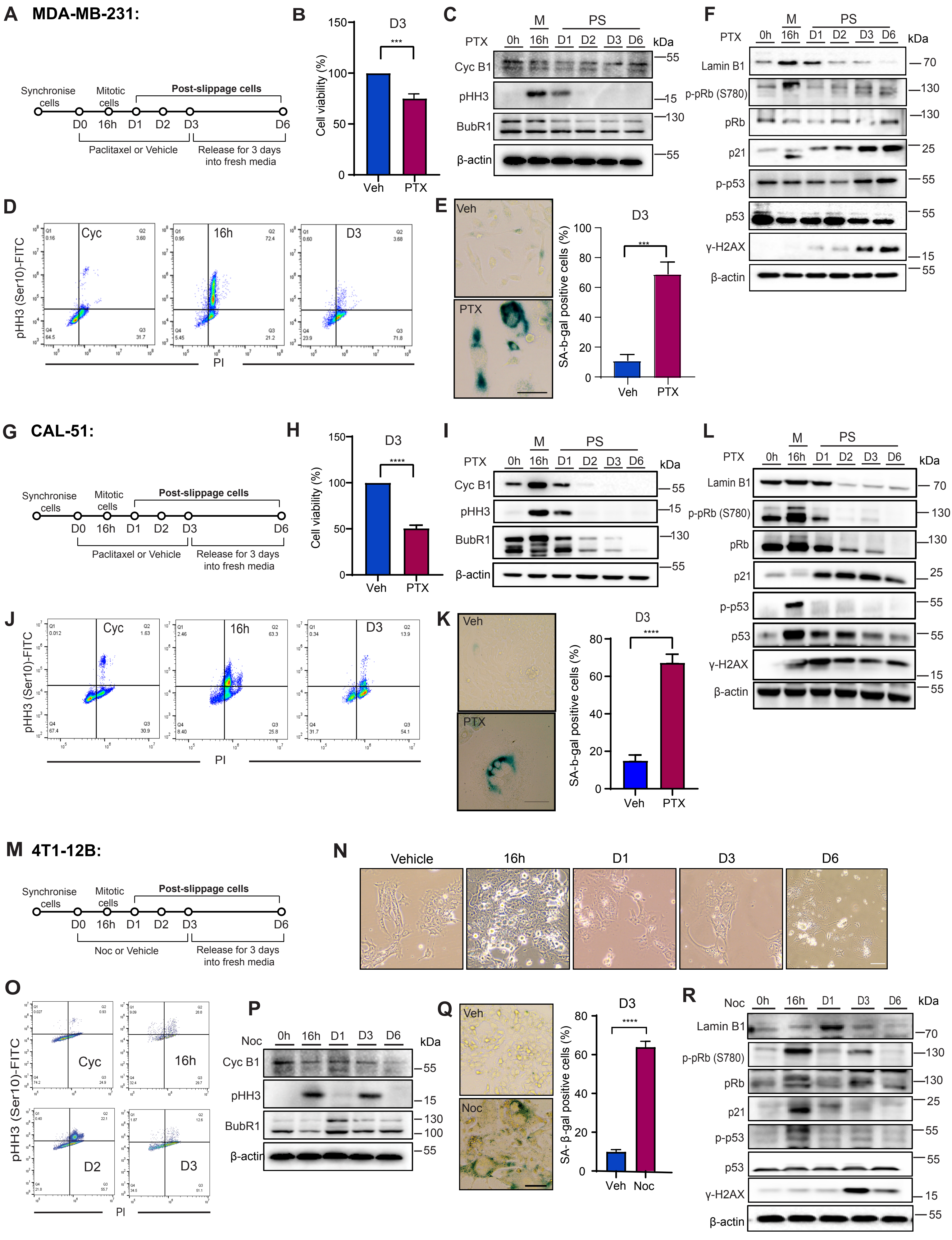
