## Supplemental Figure 2 for "Tumour-derived extracellular vesicles within the therapy-induced senescent secretome distinctly suppress breast cancer via DKK1-mediated inflammatory response"

### Supplementary Figure S2

**A**

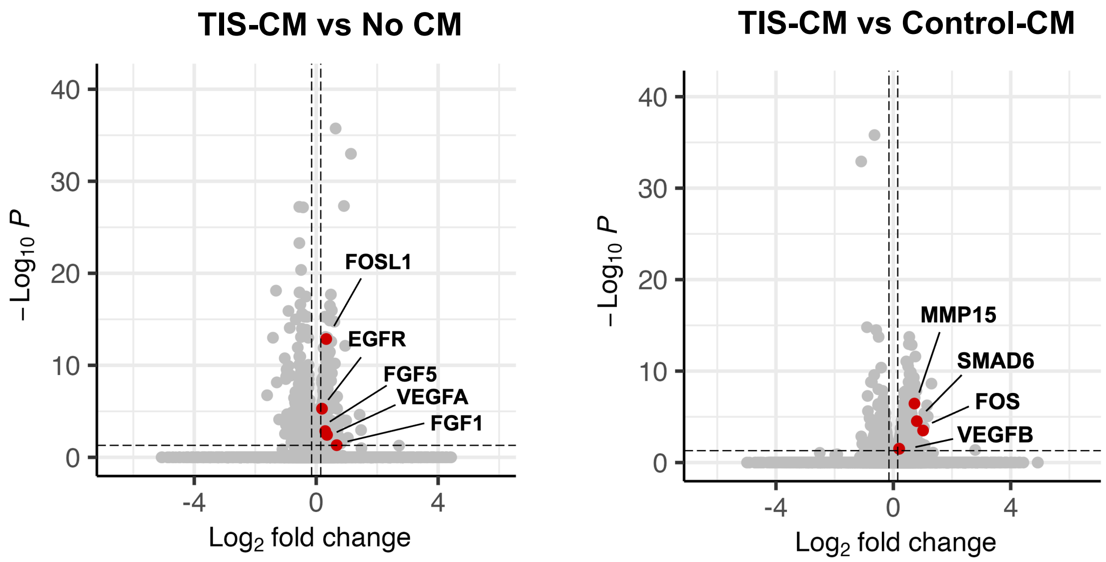


**B**

| TIS-CM vs No CM | | | | | | |
|---|---|---|---|---|---|---|
| Rank | Perturbation | Score | Mode of action | Cells | Dose | Time |
| 1 | dasatinib | 1.4315 | SRC family inhibitor | A549 | 1.11 µM | 24h |
| 2 | pelitinib | 1.4057 | EGFR inhibitor | MCF10A | 10 µM | 24h |
| 3 | radicicol | 1.3876 | HSP90 inhibitor | MCF10A | 1.11 µM | 24h |
| 4 | TWS-119 | 1.3873 | GSK-3β inhibitor | MCF10A | 10 µM | 24h |
| 5 | dasatinib | 1.383 | SRC family inhibitor | A549 | 0.04 µM | 24h |
| 6 | PD-184352 | 1.382 | MEK1/2 inhibitor | HT29 | 1.11 µM | 24h |
| 7 | dasatinib | 1.3781 | SRC family inhibitor | A549 | 3.33 µM | 24h |

| TIS-CM vs Control-CM | | | | | | |
|---|---|---|---|---|---|---|
| Rank | Perturbation | Score | Mode of action | Cells | Dose | Time |
| 1 | digoxin | 1.401 | Na/K ATPase inhibitor | MCF7 | 10 µM | 24h |
| 2 | DG-041 | 1.3962 | EP3 receptor antagonist | SW620 | 40.0 µM | 6h |
| 3 | Emetine Dihydrochloride Hydrate (74) | 1.3938 | Protein synthesis inhibitor | HT29 | 0.63 µM | 24h |
| 4 | Narciclasine | 1.3935 | Rho/ROCK/LIM kinase/cofilin modulator | HT29 | 10.0 µM | 24h |
| 5 | BRD-K56196992 | 1.3793 | - | VCAP | 10.0 µM | 24h |
| 6 | SU11652 | 1.3734 | PDGFRβ/VEGFR2/FGFR1 inhibitor | VCAP | 10.0 µM | 24h |
| 7 | BJM-ctd2-9 | 1.3706 | - | DV90 | 10.0 µM | 6h |

**Supplementary Figure S2. SASP factors derived from TIS MDA-MB-231 cells induce growth factor-related and angiogenic signalling.** (**A**) Volcano plot of DEGs in MDA-MB-231 cells incubated for 48 hours with TIS-CM vs No CM (left) and TIS-CM vs Control-CM (right) with growth factor and angiogenic signalling-associated genes highlighted. **(B)** Connectivity Map analysis showing top pertubation hits based on transcriptomic expression that could potentially reverse the tumourigenic phenotype exerted by TIS-CM.
