## Supplemental Figure 3 for "Tumour-derived extracellular vesicles within the therapy-induced senescent secretome distinctly suppress breast cancer via DKK1-mediated inflammatory response"

### Supplementary Figure S3

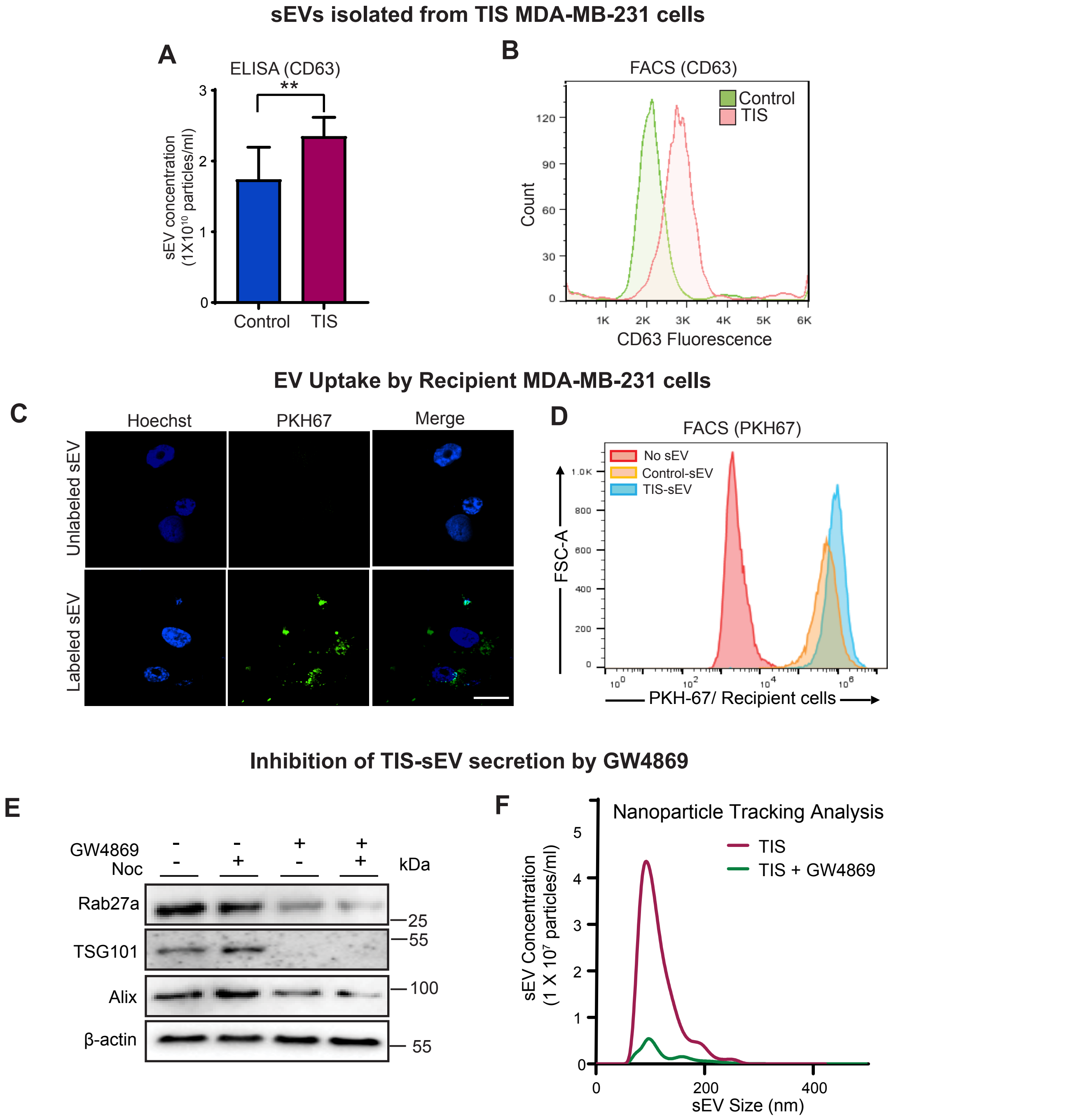


**Supplementary Figure S3. Increased sEV abundance in TIS MDA-MB-231 cells.** (**A**) sEVs derived from cells treated with DMSO (control) or nocodazole-induced senescent cells (TIS) (SEC fractions 7-12) were quantified using ELISA-based CD63. **(B)** Fractions as per (A) were incubated with CD63 antibody-coated magnetic beads and detected by flow cytometry using CD63-PE antibody. (**C**) Representative IF images of PKH67-labelled and unlabelled sEVs (2 x $10^9$ particles/ml) following uptake by recipient MDA-MB-231 cells. Scale bar: 20µm. (**D**) Flow cytometric analysis of recipient MDA-MB-231 cells incubated with PKH67-labeled sEV derived from control and TIS MDA-MB-231 cells. Data are mean ± s.d. of 3 independent experiments. **p < 0.01, ***p < 0.001 by Student's *t*-test. **(E)** Immunoblot detection of sEV marker proteins (Rab27a, TSG101, Alix) in MDA-MB-231 cells treated with 100 ng/ml of nocodazole and 15 µm of exosome inhibitor GW4869, either individually or in combination, for a duration of 72 hours. (**F**) Quantification of sEV size (nm) and number (1X10□ particles/ml) using nanoparticle tracking analysis (NTA) from TIS cells with or without treatment with GW4869.
