## Supplemental Figure 4 for "Tumour-derived extracellular vesicles within the therapy-induced senescent secretome distinctly suppress breast cancer via DKK1-mediated inflammatory response"

### Supplementary Figure S4

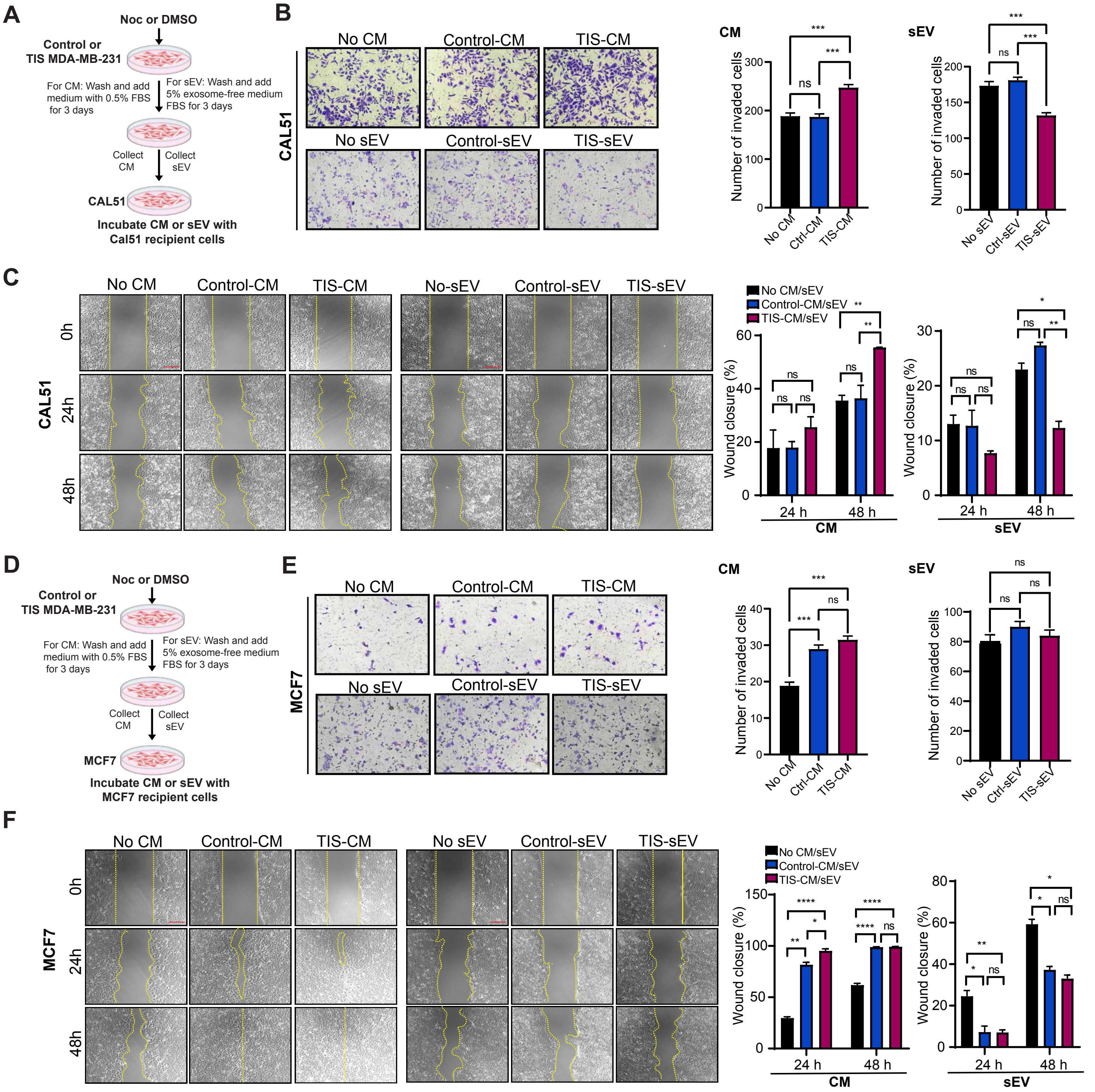


**Supplementary Figure S4. TIS cancer cell-elicited SASP and sEV confer differential tumourigenic impact on recipient cells.** (**A**) Schematic for p53 wildtype CAL51 TNBC (recipient cell) incubated with Control-CM or sEV and TIS-CM or sEV (2 x $10^9$ particles/ml) derived from MDA-MB-231 cells. **(B)** Representative images for invasion assay at 72h after start of experiment (left). Plot shows quantification of cells with invasive capabilities (right). **(C)** Representative images of wound healing assay at 48h after start of experiment. Scale bar 200 µm (left). Plot shows rate of migration (right). (**D**) Schematic for non-transformed MCF7 (recipient cell) incubated with Control-CM or sEV and TIS-CM or sEV (2 x $10^9$ particles/ml) derived from MDA-MB-231 cells. **(B)** Representative images for invasion assay at 72h after start of experiment (left). Plot shows quantification of cells with invasive capabilities (right). **(C)** Representative images of wound healing assay at 48h after start of experiment (left). Plot shows rate of migration (right). Scale bar 200 µm. Data are mean ± s.d. of 3 independent experiments; *p < 0.05, **p < 0.01, ***p < 0.001, ****p < 0.0001 and n.s. non significance by ANOVA.
