## Supplemental Figure 5 for "Tumour-derived extracellular vesicles within the therapy-induced senescent secretome distinctly suppress breast cancer via DKK1-mediated inflammatory response"

### Supplementary Figure S5

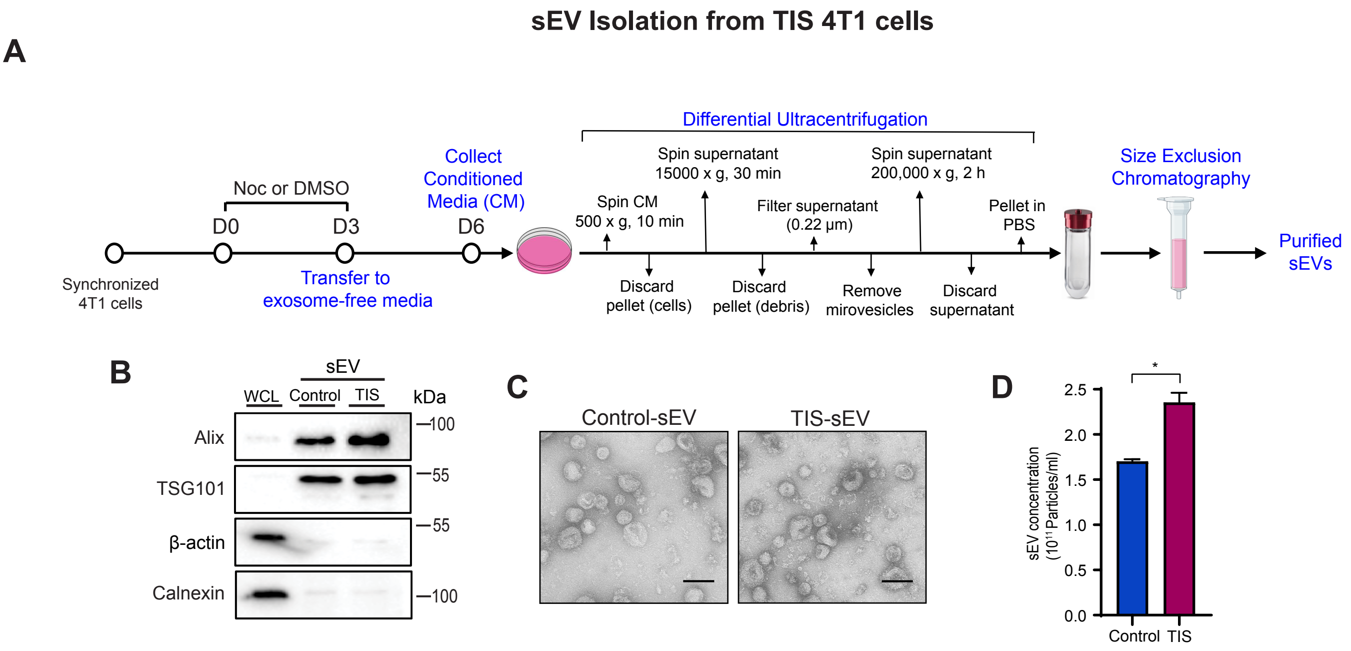


**Supplementary Figure S5. sEV isolation from Nocodazole-induced senescent murine 4T1 breast cancer cells** (**A**) Scheme for sEV isolation from control or TIS 4T1 cells using differential ultracentrifugation and size exclusion chromatography method. (**B**) Immunoblotting of whole cell lysate (WCL) and sEV-derived protein for sEV markers (Alix, TSG101) and negative controls (beta-actin, calnexin). (**C**) Representative transmission electron microscopy images of sEVs isolated from Control and TIS 4T1 cells. Scale bar, 100 nm. (**D**) sEV collected from 4T1 cells were quantified by acetylcholinesterase activity assay. Data are mean ± s.d. of 3 independent experiments; *p < 0.05 by Student's *t*-test.
