## Supplemental Figure 6 for "Tumour-derived extracellular vesicles within the therapy-induced senescent secretome distinctly suppress breast cancer via DKK1-mediated inflammatory response"

### Supplementary Figure S6

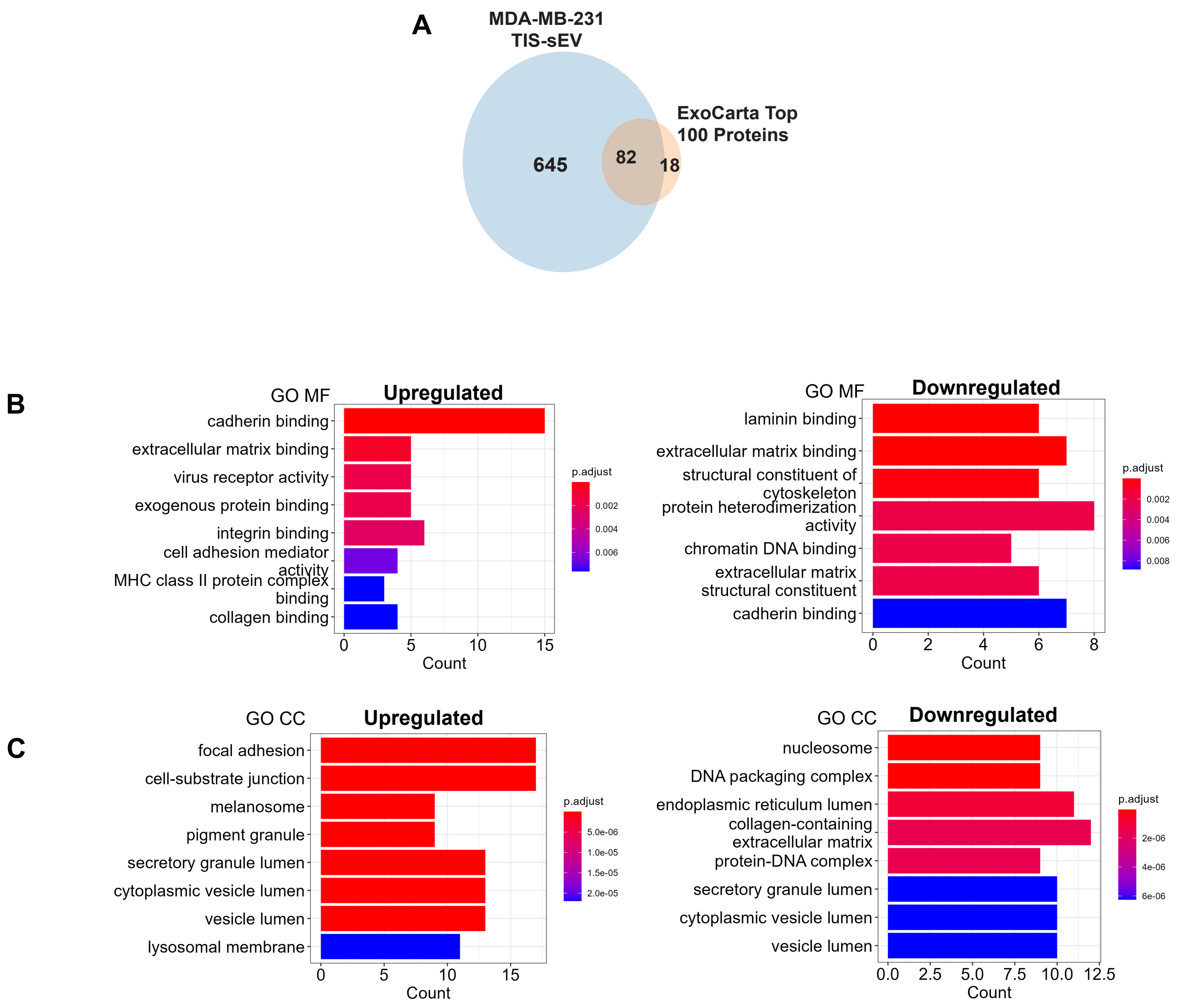


**Supplementary Figure S6. Proteins identified in TIS-sEVs are of vesicular origin.** (**A**) Venn diagram of identified proteins in TIS-sEV that overlapped with Top 100 Proteins reported on ExoCarta. (**B**) GO Molecular Function (MF) enrichment analysis of significantly upregulated and downregulated proteins found in TIS-sEV compared to Control-sEV. (**C**) GO Cellular Compartment (CC) enrichment analysis of significantly upregulated and downregulated proteins found in TIS-sEV compared to Control-sEV.
