## Supplemental Figure 7 for "Tumour-derived extracellular vesicles within the therapy-induced senescent secretome distinctly suppress breast cancer via DKK1-mediated inflammatory response"

### Supplementary Figure S7

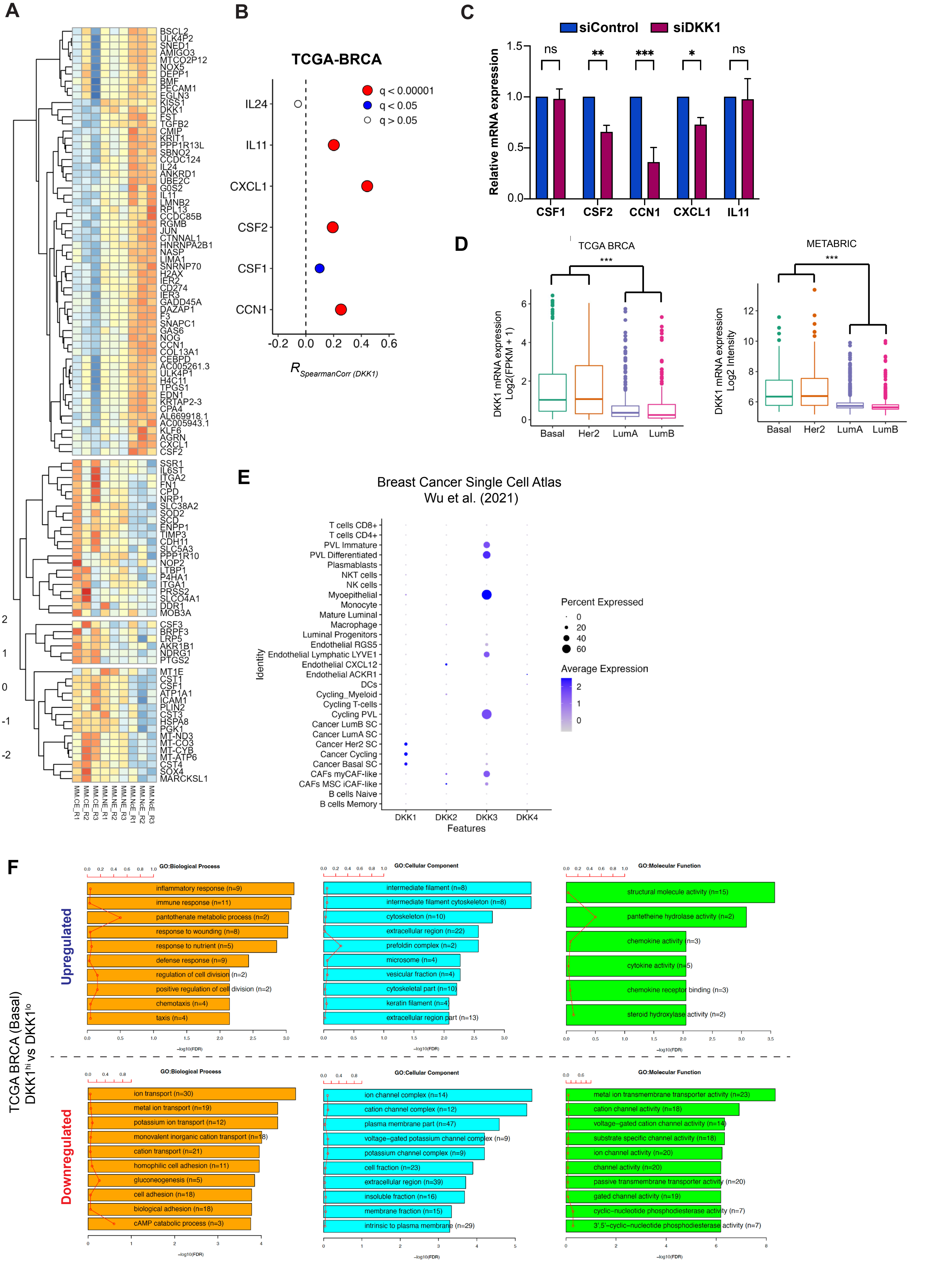

**Supplementary Figure S7. Knockdown of DKK1 downregulates CSF2, CCN1, and CXCL1 expression. (A)** Heatmap of significant differentially expressed genes as identified by RNA-seq, between cells incubated with No sEV, Control-sEV, and TIS-sEV ($2 \times 10^9$ particles/ml) for 72 hours. Right panel is snapshot of DEGs for cytokines. **(B)** Co-expression correlation analysis of DKK1 and selected cytokines in TCGA-BRCA patient cohort. **(C)** RT-PCR analysis of mRNA expression levels of CSF1, CSF2, CCN1, CXCL1, and IL11 in siControl (scrambled) vs siDKK1 cells. **(D)** DKK1 expression across breast cancer subtypes in TCGA BRCA and METABRIC cohorts. **(E)** Expression patterns of the DKK protein family across different cell types in the tumour milieu as identified in the Breast Cancer Single Cell Atlas dataset. **(F)** GO enrichment analysis of upregulated and downregulated DEGs identified in $DKK1^{hi}$ (>$75^{th}$ percentile) vs $DKK1^{lo}$ (< $25^{th}$ percentile) using TCGA BRCA basal only cohort. *$p < 0.05$, **$p < 0.01$, ***$p < 0.001$, ****$p < 0.0001$ and n.s. non significance by ANOVA. *$p < 0.05$, **$p < 0.01$, ***$p < 0.001$, ****$p < 0.0001$ and n.s. non significance by ANOVA.
